# Modulation of the *Pseudomonas aeruginosa* quorum sensing cascade by MexT-regulated factors

**DOI:** 10.1101/2025.03.17.643737

**Authors:** Andrew Frando, Robert S. Parsek, Jamal Omar, Nicole E. Smalley, Ajai A. Dandekar

## Abstract

*Pseudomonas aeruginosa (Pa)* uses quorum sensing (QS), a cell-cell communication system that enables it to sense cell density and to alter gene expression. *Pa* has three complete QS circuits controlled by the regulators LasR, RhlR, and PqsR, that together activate hundreds of genes. In the well-studied strain PAO1, QS is organized hierarchically, with PqsR and RhlR activity dependent on LasR. This hierarchy depends on the non-QS transcription factor MexT; deletion of *mexT* allows for RhlR activity in the absence of LasR. We aimed to identify how MexT modulates the *Pa* QS architecture. We compared the transcriptome of PAO1 to that of PAO1Δ*mexT* and determined a MexT regulon. We identified two MexT-regulated operons that may affect the QS hierarchy: the efflux pump genes *mexEF*-*oprN* and the *Pseudomonas* quinolone signal (PQS) synthesis genes *pqsABCDE*. We tested whether the products of these genes affected the QS hierarchy. A *mexEF* knockout mutant, like a *mexT* deletion mutant, exhibited RhlR activity earlier, and to a higher magnitude, than wild-type PAO1. MexEF-OprN is known to export quinolones, and we found that exogenous addition of PQS, through PqsE, also resulted in earlier and higher magnitude of RhlR activity compared to wild-type PAO1. We also discovered alternate QS architectures in clinical isolates, where RhlR activity is not fully dependent on LasR. In these isolates, surprisingly, MexT does not influence the relationship between LasR and RhlR. Our work reveals a new suite of factors that regulate QS in *Pa*, with implications for bacterial behaviors in environmental and clinical settings.

**Importance:** Bacteria interact with both abiotic and biotic factors in their environment. Quorum sensing (QS) is one mechanism that bacteria use to communicate with other bacteria and coordinate behaviors in the population. QS regulates a wide variety of processes ranging from the production of light to modulation of virulence factors; some bacteria use single QS circuits, while others have several. The opportunistic pathogen *Pseudomonas aeruginosa* uses QS to control some virulence functions and has three complete QS circuits. Our study explores why bacteria might have multiple QS circuits. We show how a non-QS transcription factor, MexT, influences QS regulators in *P. aeruginosa* and we uncover the diversity of QS architectures in clinical isolates. This study begins to reveal the benefits (or disadvantages) of multiple QS circuits, allowing us to understand behaviors of bacteria that have a range of implications including in health, agriculture, and bioremediation.

## Introduction

Bacteria can occupy a variety of environments that include the ocean, human body, and soil, and seldom live in isolation. Bacteria have many methods of adapting to environmental changes and interacting with other microbes, including by using various forms of signaling. Quorum sensing (QS) is a type of intercellular communication used by many bacteria to coordinate group behaviors in response to cell density. QS is a feature of many gram-positive and gram-negative bacteria and typically results in transcription factor activation of gene expression (1). Bacterial behaviors regulated by QS include the production of light, biofilm formation, and the secretion of virulence factors.

QS is typically organized into circuits composed of a signal producing enzyme and a signal-responsive transcriptional regulator. *Pseudomonas aeruginosa* (*Pa*) is a gram-negative opportunistic bacterial pathogen that encodes multiple QS circuits that are controlled by the transcriptional regulators LasR, RhlR, and PqsR (also called MvfR) (2). Together, the three QS circuits regulate 5% of gene expression, including virulence factors like proteases, toxins, and biofilm components (3).

LasR and RhlR are both QS transcription factors that respond to acyl-homoserine lactone (AHL) signals (4). LasI produces the signal *N*-3-oxo-dodecanoyl homoserine lactone (3OC12-HSL) that binds to and results in LasR activity. Similarly, RhlI produces the signal *N-*butanoyl homoserine lactone (C4-HSL) that binds to RhlR. The PqsR QS circuit involves the quinolone signals 2-heptyl-4-quinolone (HHQ) and 2-heptyl-3-hydroxy-4(1H)-quinolone (PQS) (5–8). In a process different from AHL QS, HHQ and PQS are synthesized from multiple enzymes encoded by *pqsABCD* and *pqsH*. All three circuits behave in a positive feedback loop where signals that bind activate the transcriptional regulator, which in turn results in increased expression of the signal synthase enzymes (9), a typical feature of QS circuits.

The interconnectivity between the QS circuits, or QS architecture, differs between *Pa* strains. In the laboratory strain PAO1, QS architecture is hierarchical, where PqsR and RhlR expression are dependent on LasR (10). A *lasR*-null strain in PAO1 is generally in a QS-off state (11). There are additional layers of interaction: the chaperone PqsE, which is regulated by PQS QS, has been shown to positively affect RhlR activity, while RhlR negatively regulates production of PQS (12–14). The QS architecture in some clinical isolates seems to vary from PAO1. We and others have found that despite encoding inactivating mutations in *lasR*, *Pa* clinical isolates derived from chronic infections of people with cystic fibrosis are capable of activating RhlR QS independent of LasR, in a manner that differs from PAO1 (15–17). The question remains whether these clinical isolates have QS architectures that are fundamentally different from PAO1 or if the phenotype is explained, instead, by mutations that were acquired in addition to those in *lasR*.

More recent work has identified how PAO1 can itself develop a LasR-independent QS architecture. The “wildtype” strain PAO1 acquired a loss-of-function mutation in the oxidoreductase gene *mexS*, presumably as a consequence of exposure to chloramphenicol (18–20). MexS responds to oxidative stress and negatively regulates the activity of a transcriptional regulator, MexT. Because of the inactivating mutation in *mexS,* MexT is constitutively active in PAO1 and induces many genes including the RND drug efflux pump MexEF-OprN. MexEF-OprN has been shown to influence QS, in part by delaying the activation of PQS QS (21). Recent work has linked MexT to the regulation of QS in PAO1 (22–24). PAO1 with inactivating mutations in *lasR* do not activate RhlR QS. However, *lasR* mutants exhibit RhlR activity if there is also an inactivating mutation in the *mexT*. These data indicate that MexT contributes to the LasR-dependent QS architecture in PAO1, but the mechanisms driving this phenomenon are unknown.

We are interested in QS architectures and how they are regulated, with the idea that the arrangement of QS circuits might impact bacterial virulence, social behaviors, or both. To investigate this issue, we explored how MexT regulates QS in the laboratory strain PAO1. We determined the MexT regulon by comparing the transcriptomes of wild-type PAO1 to a *mexT*-null mutant. We ascertained that the efflux pump MexEF-OprN and the chaperone protein PqsE both contribute to the LasR-dependent QS architecture in PAO1. To study the diversity of quorum sensing architectures, we next explored QS architecture in clinical isolates with functional LasR alleles. We identified isolates with a PAO1-like QS architecture where RhlR activity is dependent, to varying degrees, on intact LasR, expanding the array of known QS architectures in *Pa*. Finally, we determined that MexT did not influence QS architecture in some of these clinical isolates, suggesting the existence of other regulators of the QS hierarchy in *Pa*.

## Results

### The MexT regulon

Because inactivating mutations in *mexT* alter the QS hierarchy in PAO1, we started by exploring how MexT regulates quorum sensing architecture in this strain. MexT is a transcription factor that is known to regulate several genes (25, 26), although the full extent of its gene regulation is not fully known, because a prior study compared the wild-type to an overexpression variant (26), and we now understand that MexT is constitutively expressed in PAO1. We aimed to identify MexT-regulated genes that modulate QS, specifically components of the MexT regulon that might impact the timing of RhlR QS.

We compared RhlR activity in a PAO1 *mexT* knockout mutant (PAO1Δ*mexT*), to that of wild-type PAO1 (**Figure 1**). In agreement with prior studies (22, 23), we found that the PAO1Δ*mexT* mutant exhibited higher RhlR activity at lower cell densities compared to wild-type. The PAO1Δ*mexT* knockout mutant has robust RhlR activity as early as an OD_600_ of 1.0, while wild-type PAO1 exhibits RhlR activity around an OD_600_ of 2.0. We identified the OD_600_ of 1.0 and 2.0 as the specific growth stages we would use to identify MexT-regulated genes that might modulate RhlR QS.

**Figure 1.**
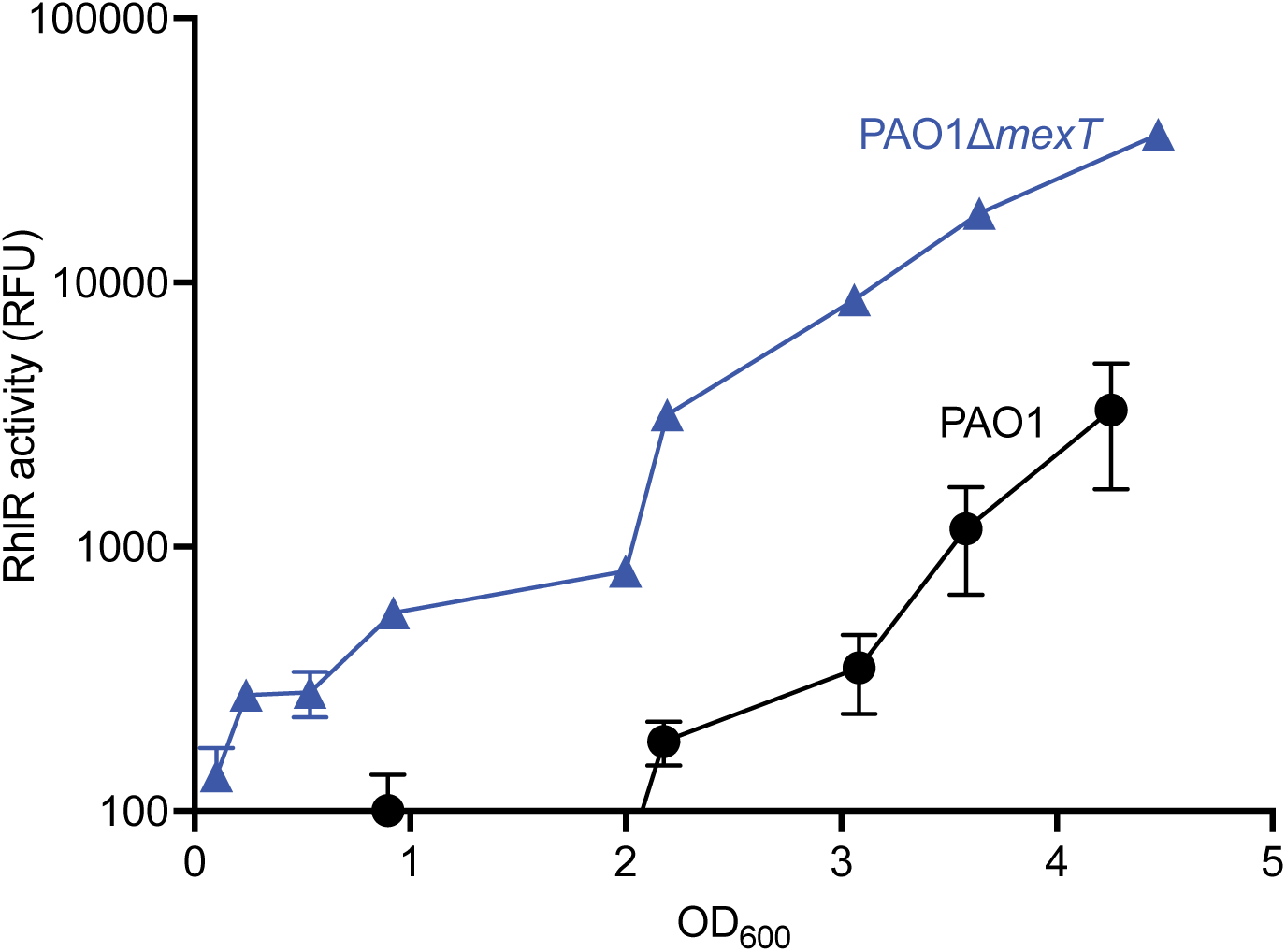
PAO1Δ*mexT* exhibits earlier and greater RhlR activity than the wild-type. Wild-type PAO1 and PAO1Δ*mexT* were grown in flasks for 4.5 hours. Bacterial growth was determined by OD_600_, and RhlR activity was determined using a reporter plasmid with the promoter of the RhlR-regulated gene *rhlA* controlling GFP expression. The x-axis shows OD_600_, and the y-axis shows RhlR activity as measured using the *rhlA-gfp* reporter plasmid.

We performed an RNA-sequencing experiment using wild-type PAO1 and a *mexT* knockout mutant to identify differentially expressed genes (DEGs) at OD_600_ of 1.0 and 2.0. We compared gene expression between the two strains using DESeq2 (27) and determined the MexT regulon. We defined DEGs as genes in the *mexT* knockout mutant that have at least a 2-fold change compared to wild-type PAO1 and a *p-*value < 0.05. In total, we identified 152 DEGs at an OD_600_ of 1.0, with 104 genes exhibiting higher expression in the *mexT* knockout mutant and 48 genes having higher expression in wild-type PAO1 (**Supplemental Table 1**). We identified 265 DEGs at an OD_600_ of 2.0, with 178 genes showing higher expression in the *mexT* knockout mutant and 87 genes having higher expression in wild-type PAO1 (**Supplemental Table 2**). Based on this experiment, the MexT regulon increases during growth. We observed 98 genes shared between the regulon at an OD_600_ of 1.0 and 2.0, demonstrating that over 60% of genes at OD_600_ of 1.0 continued to be expressed and differentially regulated later in growth.

We observed that a PAO1Δ*mexT* mutant showed an earlier and higher magnitude of RhlR activation, indicating that MexT likely negatively regulates RhlR in wild-type PAO1 (**Figure 1**). Consistent with this observation, the transcriptomes we identified include several genes regulated by RhlR. These genes are probably not contributing to MexT-specific regulation of QS. Therefore, we filtered out genes that belong to the core RhlR regulon that was identified previously using the laboratory strain PAO1 and clinical isolates derived from people with cystic fibrosis (**Figure 2 and Table 1)** (15, 16, 28). Because there was robust RhlR activation at both OD_600_ of 1.0 and 2.0, we focused on the OD_600_ 1.0 transcriptome for the remainder of our work, since it was more likely to include factors relevant to RhlR regulation and fewer genes that are activated by RhlR itself.

**Figure 2.**
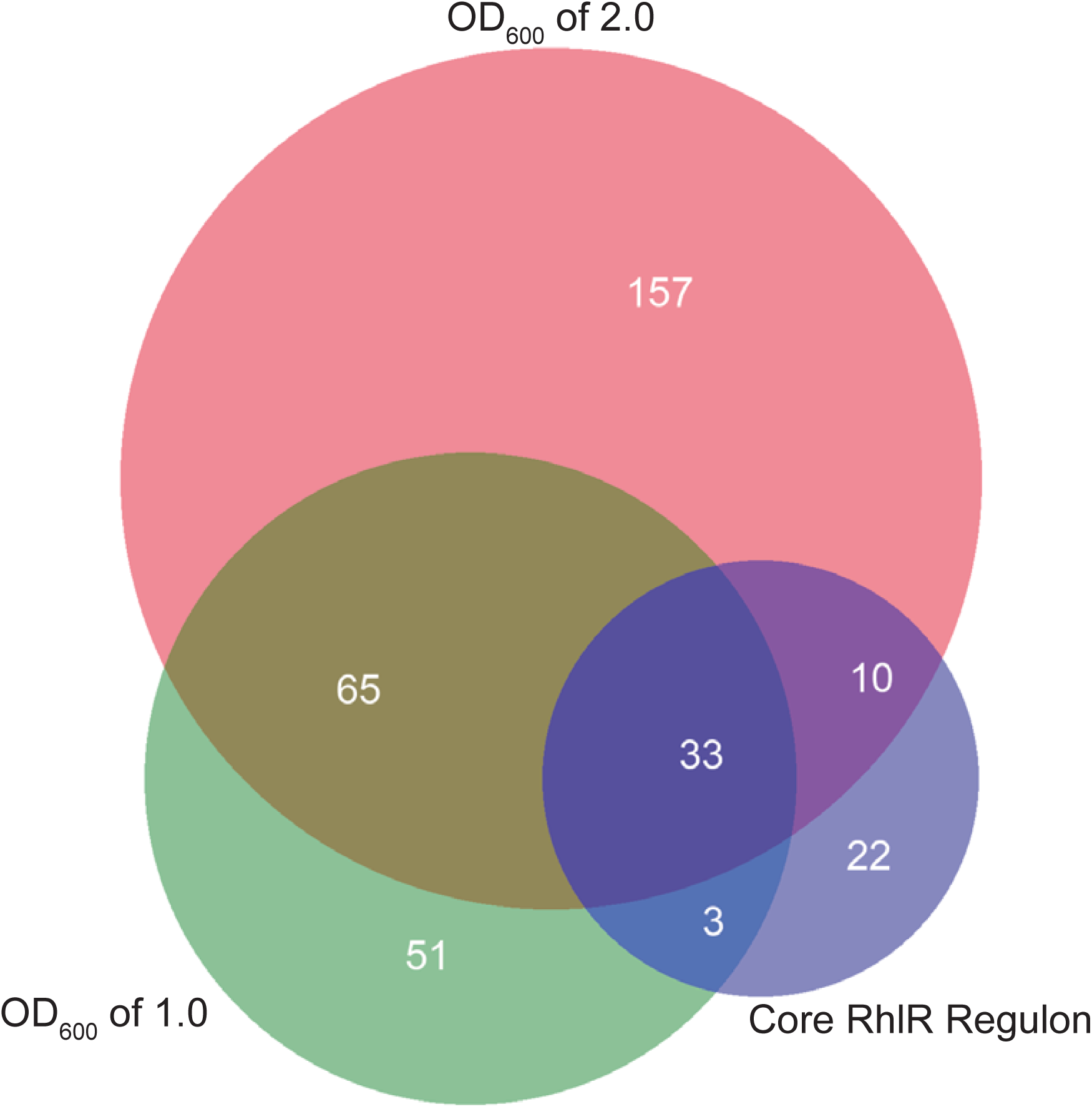
The MexT regulon encompasses at least 150 genes and overlaps with the RhlR regulon. Venn diagram showing the overlap of genes between an OD_600_ of 1.0 and 2.0, and a previously identified RhlR regulon (15, 16, 28).

**Table 1.** The MexT regulon. . We performed RNA-seq on wild-type PAO1 and PAO1Δ*mexT* grown to an OD_600_ of 1.0. Differentially expressed genes are defined as genes that showed a 2-fold change compared to wild-type PAO1 and had a *p*-value less than 0.05. We further filtered the differentially expressed genes to exclude those that belong to the core RhlR regulon.

| Gene expression changes in PAO1ΔmexT compared to PAO1 | | | | | | | | |
| --- | --- | --- | --- | --- | --- | --- | --- | --- |
| Gene | log<sub>2</sub> fold-change | p-value | Gene | log<sub>2</sub> fold-change | p-value | Gene | log<sub>2</sub> fold-change | p-value |
| PA4881 | -6.5 | 6.71E-30 | PA2811 | -1.3 | 9.80E-17 | opdC | 1.3 | 4.27E-16 |
| mexF | -6.4 | <1.0E-40 | PA2358 | -1.3 | 2.19E-02 | opmD | 1.3 | 1.46E-04 |
| mexE | -6.3 | <1.0E-40 | pagL | -1.2 | 1.78E-20 | kdpB | 1.3 | 1.44E-05 |
| PA3229 | -6.1 | <1.0E-40 | PA2812 | -1.2 | 3.53E-07 | pqsA | 1.4 | 3.63E-15 |
| PA1942 | -6.0 | <1.0E-40 | nosL | -1.2 | 6.82E-04 | stp1 | 1.4 | 3.19E-06 |
| oprN | -6.0 | <1.0E-40 | PA3283 | -1.1 | 9.10E-05 | pqsC | 1.4 | 1.19E-05 |
| PA4623 | -5.8 | <1.0E-40 | PA2354 | -1.1 | 1.65E-03 | mexH | 1.4 | 3.65E-07 |
| PA1744 | -4.6 | <1.0E-40 | bexR | -1.1 | 4.11E-12 | ladS | 1.4 | 5.18E-26 |
| PA2486 | -4.5 | <1.0E-40 | PA1418 | -1.1 | 1.70E-02 | pqsB | 1.4 | 1.64E-07 |
| PA1743 | -4.2 | <1.0E-40 | chiC | 1.0 | 1.19E-03 | pscF | 1.4 | 1.05E-03 |
| PA3871 | -3.8 | 6.57E-05 | PA1658 | 1.0 | 3.37E-02 | PA1669 | 1.4 | 4.88E-11 |
| PA4354 | -3.8 | <1.0E-40 | pchA | 1.0 | 8.82E-04 | PA1668 | 1.4 | 8.81E-07 |
| narI | -3.8 | 5.45E-05 | oprB | 1.0 | 1.77E-08 | pchE | 1.4 | 4.11E-12 |
| PA2487 | -3.7 | <1.0E-40 | phzS | 1.0 | 5.93E-04 | phzD2 | 1.4 | 5.31E-04 |
| narJ | -3.7 | 3.32E-04 | PA2097 | 1.0 | 1.21E-02 | phnA | 1.4 | 7.42E-10 |
| moaA1 | -3.6 | 4.17E-05 | PA2362 | 1.1 | 3.20E-02 | rbsC | 1.5 | 3.19E-06 |
| PA2485 | -3.5 | <1.0E-40 | PA3377 | 1.1 | 2.71E-03 | PA2782 | 1.5 | 4.88E-05 |
| PA1970 | -3.5 | <1.0E-40 | PA4341 | 1.1 | 3.22E-02 | PA2068 | 1.5 | 1.44E-05 |
| PA3230 | -3.2 | <1.0E-40 | phzE2 | 1.1 | 3.10E-04 | PA3574a | 1.5 | 5.04E-03 |
| PA4355 | -2.9 | 1.10E-29 | bkdA1 | 1.1 | 1.50E-02 | PA2067 | 1.6 | 3.12E-03 |
| narH | -2.7 | 2.04E-05 | pscE | 1.1 | 7.13E-03 | hcpA | 1.6 | 5.93E-06 |
| gbuA | -2.6 | 2.06E-14 | PA4219 | 1.1 | 1.01E-03 | PA2781 | 1.6 | 4.08E-08 |
| PA1333 | -2.6 | 1.20E-24 | kdpC | 1.1 | 3.09E-02 | rbsA | 1.6 | 1.21E-03 |
| PA2813 | -2.5 | <1.0E-40 | PA3677 | 1.1 | 1.19E-03 | PA4142 | 1.7 | 3.10E-14 |
| xenB | -2.4 | 3.19E-21 | stk1 | 1.2 | 1.09E-06 | hcpB | 1.7 | 5.57E-06 |
| nosY | -2.2 | 2.17E-04 | PA1657 | 1.2 | 5.06E-03 | pchF | 1.7 | 5.28E-04 |
| PA2491 | -2.2 | <1.0E-40 | pchD | 1.2 | 4.78E-04 | PA2260 | 1.8 | 1.49E-07 |
| PA1419 | -2.1 | 1.04E-09 | phzF2 | 1.2 | 1.57E-03 | pchC | 1.8 | 1.49E-07 |
| nirN | -2.1 | 5.30E-06 | pqsD | 1.2 | 1.05E-04 | hisJ | 1.8 | 1.55E-02 |
| PA0510 | -2.1 | 1.17E-08 | hcpC | 1.2 | 1.42E-03 | pchB | 1.8 | 1.48E-07 |
| PA2759 | -2.0 | 2.58E-30 | PA0050 | 1.2 | 6.64E-05 | PA3519 | 1.8 | 7.49E-03 |
| PA4882 | -2.0 | 7.54E-35 | PA2322 | 1.2 | 1.09E-03 | PA4596 | 1.8 | 1.56E-02 |
| PA1420 | -1.9 | 1.30E-14 | popN | 1.3 | 3.56E-04 | PA2069 | 1.8 | 1.18E-11 |
| PA3381 | -1.7 | 5.26E-03 | bkdA2 | 1.3 | 5.90E-10 | oprD | 1.9 | 2.63E-27 |
| PA1417 | -1.6 | 3.35E-06 | hpcG | 1.3 | 1.58E-12 | PA3325 | 2.0 | 1.37E-25 |
| PA2758 | -1.6 | 5.48E-22 | PA5180 | 1.3 | 1.19E-03 | PA5328 | 2.2 | 1.60E-02 |
| PA3282 | -1.5 | 3.62E-04 | pqsE | 1.3 | 1.19E-05 | pcrG | 2.2 | 1.28E-04 |
| PA1416 | -1.5 | 5.77E-06 | PA4218 | 1.3 | 2.23E-05 | PA0165 | 2.3 | 7.90E-33 |
|  |  |  |  |  |  | PA4133 | 2.7 | 1.12E-27 |

Using the OD_600_ 1.0 transcriptome, we identified 115 genes that belong solely to the MexT regulon with 47 genes that appear to be activated by MexT (genes with lower expression in PAO1Δ*mexT* compared to wild-type PAO1) and 68 genes that appear to be repressed (genes with higher expression in PAO1Δ*mexT* compared to wild-type PAO1). Within the MexT regulon, we identified 51 genes that were identified in the previous study that used a MexT overexpression mutant (25). In total, over 30% of the genes identified in the prior study were identified in ours. The genes we identified in the MexT regulon at an OD_600_ of 1.0 belong to several protein families, including transporters, phenazine biosynthesis, and transcriptional regulators.

To identify MexT effectors that might modulate QS, we filtered for genes that have a high confidence, such as those exhibiting large fold-changes, identification of whole operons, and genes with known QS interactions. Our data validate other studies showing that MexT is a transcriptional regulator of the multidrug RND efflux pump MexEF-OprN (26, 29). MexEF-OprN is known for transport of the quorum sensing signal HHQ, a PQS precursor molecule, resulting in delayed PQS QS (21). A recent study screening for QS-inhibitory compounds identified Z-ethylthio enynone as a molecule that targets MexEF-OprN and affects C4-HSL concentrations intra- and extracellularly in wild type PAO1 (30). Given these facts, it is plausible that MexEF-OprN expression is induced by constitutive MexT activity and modulates quorum sensing architecture by altering QS signal concentrations and affecting QS activation.

### MexT regulation of efflux pumps affects the QS hierarchy

We explored the role of MexEF-OprN in modulating QS in PAO1. We hypothesized that, like MexT, MexEF-OprN negatively regulates RhlR activity. To test this, we first made a *mexEF* knockout mutant to disrupt the efflux activity of MexEF-OprN. Like other RND systems found in *P. aeruginosa*, membrane transport of substrates requires all three proteins to function (31). We then introduced episomal plasmids with reporters for LasR (*lasI-gfp*), RhlR (*rhlA-gfp*), or PqsR (*pqsA-gfp*) activity. We engineered a *mexEF* knockout mutant, which is sufficient to disrupt the efflux pump since it needs all three proteins to function, and introduced these QS reporter plasmids. The *mexEF* knockout mutant shows similar LasR activity and timing compared to wild-type PAO1 and the *mexT* knockout mutant (**Figure 3A**). However, RhlR activity was different: we observed that the *mexEF* knockout mutant, like PAO1Δ*mexT*, showed earlier and higher RhlR activity compared to wild-type PAO1 (**Figure 3B**). This finding demonstrates that the efflux pump MexEF-OprN is an important MexT-regulated effector modulating QS. Further, we found PqsR activity earlier in both the *mexT* and *mexEF* knockout mutants. In parallel, we tested the role of MexEF-OprN in regulating RhlR QS by creating an integrating plasmid that contains *mexEF-oprN* under the control of an arabinose-inducible promoter (PAO1 + *mexEF-oprN* OE) and compared this mutant to wild-type and PAO1Δ*mexT* (**Figure 3D**). When we induced expression of *mexEF-oprN,* we found that the pump alters the magnitude but not the timing of RhlR activation compared to PAO1 and PAO1 *mexEF-oprN* OE without arabinose. These results corroborate the idea that MexEF-OprN impacts RhlR QS. Together, they demonstrate that both *mexT* and *mexEF* knockout mutants affect RhlR and PqsR QS. Because PAO1Δ*mexEF* phenocopies PAO1Δ*mexT*, it shows that MexT regulation of RhlR QS acts at least in part through the efflux pump MexEF-OprN, indicating a regulatory pathway from MexT to MexEF-OprN to RhlR QS.

**Figure 3.**
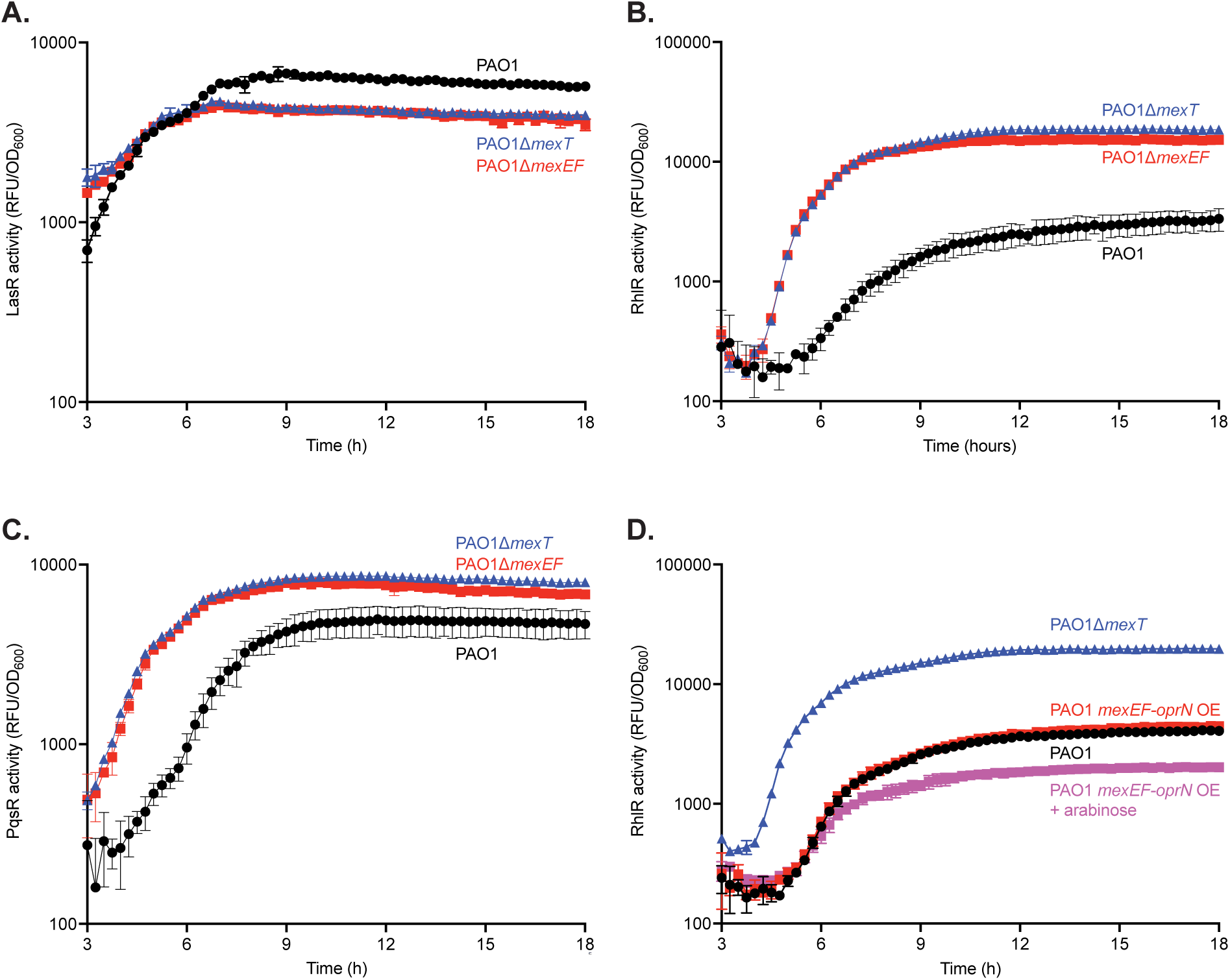
MexEF-OprN regulates both RhlR and PqsR QS. **A**. LasR activity of PAO1, PAO1Δ*mexT*, and PAO1Δ*mexEF* were determined using a LasR activity (*lasI-gfp*) reporter plasmid. **B**. RhlR activity for PAO1, PAO1Δ*mexT*, and PAO1Δ*mexEF* were determined using a RhlR activity (*rhlA-gfp*) reporter plasmid. **C**. PqsR activity for PAO1, PAO1Δ*mexT*, and PAO1Δ*mexEF* were determined using a PqsR activity (*pqsA-gfp*) reporter plasmid. **D.** RhlR activity for PAO1, PAO1Δ*mexT*, PAO1Δ*mexEF*, and PAO1 + *mexEF-oprN* OE were determined using a RhlR activity reporter plasmid.

While RhlR activity is negatively regulated by MexEF-OprN and MexT in wild-type PAO1, RhlR activity eventually occurs during later growth phases, indicating that either the negative regulation is relaxed or a positive regulatory pathway is induced. To determine whether *mexEF-oprN* expression decreases over growth and leads to the increase in RhlR activity, we performed a quantitative real-time PCR experiment to monitor *mexE* expression over bacterial growth, with each subsequent timepoint compared to time 0 (**Figure 4A**). We observed that *mexE* expression slightly increased at an OD_600_ of 1.0 and then further increased by more than 10-fold at an OD_600_ of 2.0. These results show that the increase in RhlR activity in PAO1 at later time points is not driven by a reduction in *mexEF-oprN* expression.

**Figure 4.**
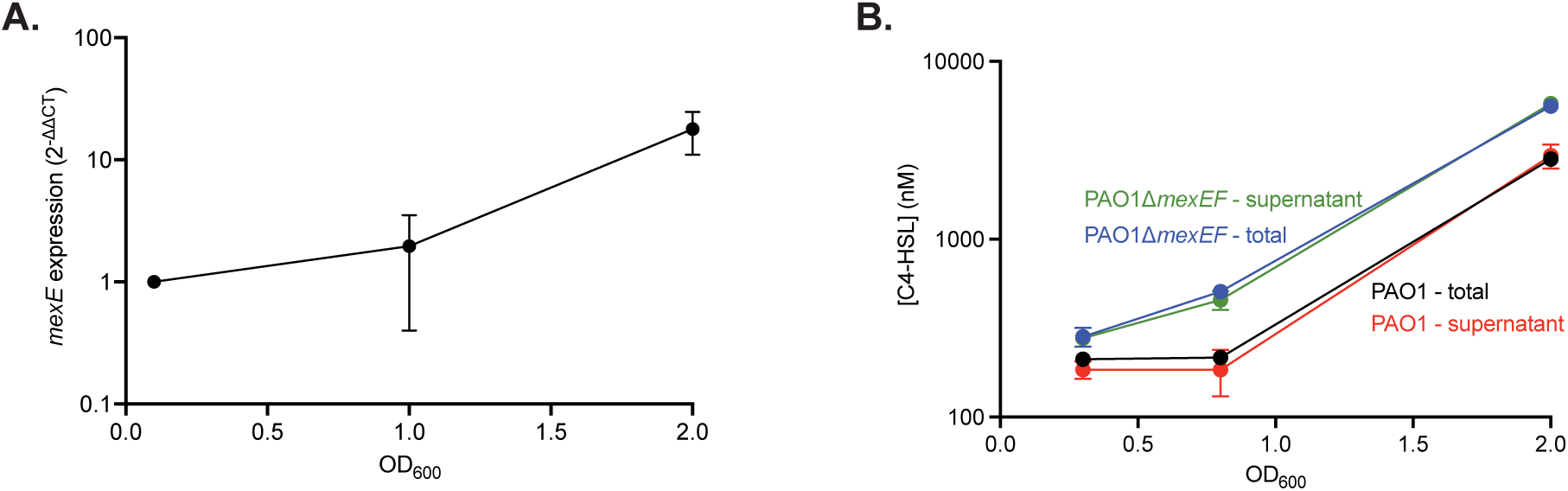
Expression of *mexEF-oprN* and C4-HSL concentrations do not account for changes in RhlR activity. **A**. Quantitative real-time PCR measuring *mexE* transcripts during bacterial growth of wild-type PAO1. *mexE* expression was normalized to *rplU* and reported as 2^-ΔΔCT^. **B**. C4-HSL was extracted from total cultures and culture supernatant over bacterial growth. C4-HSL concentrations were determined using a previously described *E. coli* reporter.

We also tested whether the delay in RhlR activity in wild-type PAO1 was due to MexEF-OprN modulating production of the RhlR signal C4-HSL. If C4-HSL is a substrate of MexEF-OprN, and MexEF-OprN delayed RhlR QS by exporting C4-HSL signal, then we would expect a higher extracellular C4-HSL concentration in wild-type PAO1 than in a *mexEF* deletion mutant. We tested this hypothesis by measuring C4-HSL concentrations in wild-type PAO1 and PAO1Δ*mexEF* over time (**Figure 4B**). We observed that total (cellular + supernatant) and extracellular (supernatant only) C4-HSL concentrations were similar between wild-type PAO1 and PAO1Δ*mexEF* at OD_600_ of 0.3. At an OD_600_ of 0.8 and 2.0, we observed activation of RhlR in the PAO1Δ*mexEF* mutant, evidenced by the increase in both total and extracellular C4-HSL concentrations. (**Figure 2**). Importantly, we do not observe any differences between total and extracellular C4-HSL concentrations in wild-type PAO1 or PAO1Δ*mexEF* at any point in growth, indicating that MexEF-OprN is not altering C4-HSL concentrations in a way that could alter RhlR activity.

### MexT regulation of PqsE also contributes to the QS hierarchy

MexEF-OprN has been shown to delay PQS QS through binding and exporting the PQS precursor HHQ (21). The gene product of *pqsE*, which is regulated by PQS QS, is known to interface with Rhl QS by positively regulating activation of some genes in the RhlR regulon (32, 33). We observed that the MexT regulon at an OD_600_ of 1.0 included the entire PQS biosynthesis operon *pqsABCDE,* which includes the gene encoding the chaperone PqsE. Because we identified these PQS biosynthesis genes in our RNA-seq experiment and have shown a role of MexEF-OprN in regulating RhlR QS, we hypothesized that constitutive MexT activity induces *mexEF-oprN* expression, leading to repression of PQS QS, delayed *pqsE* expression, and thus delaying RhlR activation.

We tested whether MexT delays PQS QS through the MexEF-OprN efflux pump. We measured PqsR activity in wild-type PAO1 supplemented with exogenous PQS and compared that to the PAO1Δ*mexT* and PAO1Δ*mexEF* mutants (**Figure 5A**). We found that addition of exogenous PQS to the wild-type culture phenocopies the PqsR activity we observed in both PAO1Δ*mexT* and PAO1Δ*mexEF*, demonstrating that the delay in PQS activity in wild-type PAO1 can be attributed to the regulatory activity of MexT and MexEF-OprN.

**Figure 5.**
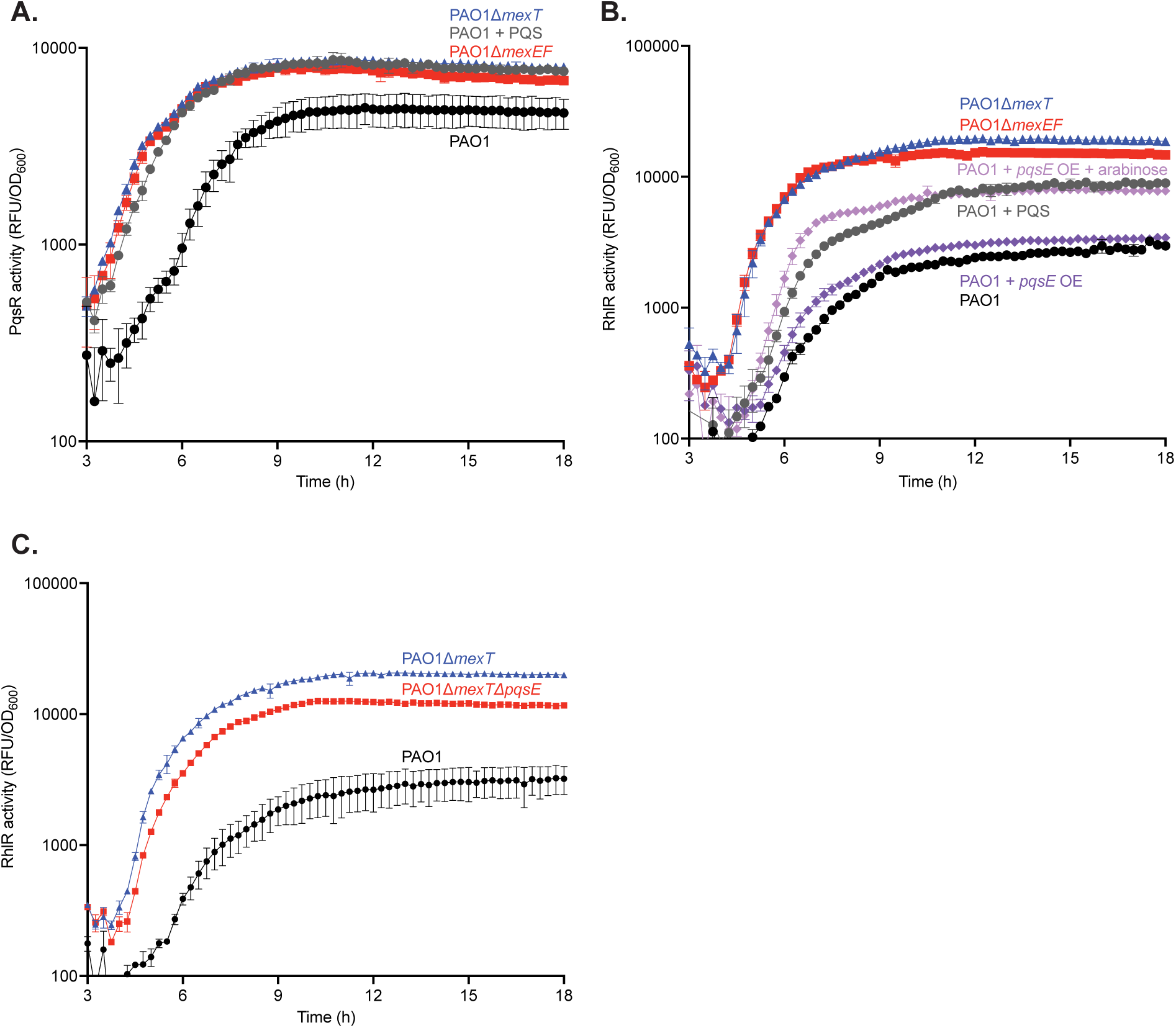
PqsE partially regulates RhlR activity. **A**. PqsR activity with and without addition of 20 µM PQS for PAO1, PAO1Δ*mexT*, and PAO1Δ*mexEF* was determined using a PqsR activity reporter plasmid. **B**. RhlR activity with and without addition of 20 µM PQS for PAO1, PAO1Δ*mexT*, PAO1Δ*mexEF*, and PAO1 + *pqsE* with and without arabinose were determined using a RhlR activity reporter plasmid. **C**. RhlR activity for PAO1, PAO1Δ*mexT*, PAO1Δ*mexT*Δ*pqsE* were determined using a RhlR activity reporter plasmid.

We next determined whether this delay in PQS QS affected RhlR QS. We measured RhlR activity in PAO1Δ*mexT* and PAO1Δ*mexEF*, and we reasoned that addition of exogenous PQS may alter RhlR activity to match that of the PAO1Δ*mexT* and PAO1Δ*mexEF* mutants, like PqsR activity (**Figure 5B**). We found that the addition of exogenous PQS did indeed result in earlier and higher RhlR activity compared to wild-type PAO1. However, the activity was intermediate between wild-type PAO1 and both PAO1Δ*mexT* and PAO1Δ*mexEF*, indicating that the delay in PQS QS only partially accounts for the relative delay of RhlR QS in wild-type PAO1.

Because we observed a change in RhlR activity with the addition of PQS, we sought to identify the factor mediating this change. PqsE is known to act as a chaperone and activate RhlR. Further, based on our transcriptome data, we observed higher expression of *pqsE* in the PAO1Δ*mexT* mutant compared to wild-type PAO1. Together, these data led us to speculate that the regulation of RhlR activity in wild-type PAO1 was due to repression of *pqsE*. To test this hypothesis, we constructed a mutant to counter *pqsE* repression mediated by MexT. This mutant contains an integrated plasmid with *pqsE* under the control of an arabinose-inducible promoter (PAO1 + *pqsE* OE), and we compared RhlR activity in this mutant to wild-type PAO1, PAO1Δ*mexT*, and PAO1Δ*mexEF* (**Figure 5B**). We found that PAO1 + *pqsE* OE without arabinose showed similar RhlR activity as wild-type PAO1, indicating negligible basal expression from the promoter. However, with the addition of arabinose, PAO1 + *pqsE* OE showed nearly identical RhlR activity to wild-type supplemented with exogenous PQS.

We further assessed the role of PqsE by creating a *mexT* and *pqsE* double knockout mutant (PAO1Δ*mexT*Δ*pqsE*) and compared RhlR activity in this mutant to wild-type PAO1 and PAO1Δ*mexT* (**Figure 5C**). In agreement with the prior experiment, we observed a decrease in RhlR activity that occurred at a slightly later timepoint in PAO1Δ*mexT*Δ*pqsE* compared to PAO1Δ*mexT*. However, PAO1Δ*mexT*Δ*pqsE* retained higher RhlR activity compared to PAO1, indicating that *pqsE* is necessary but not sufficient for MexT- and MexEF-OprN-mediated regulation of RhlR activity. These experiments showed that the increase in RhlR activity due to the addition of PQS is likely mediated by the earlier activation of PQS QS and increased production of PqsE.

Together, these data enhance prior work on MexT, MexEF-OprN, and RhlR QS (**Figure 6**). We showed that constitutive MexT activity in PAO1 resulted in increased expression of efflux pump genes *mexEF-oprN.* At lower cell densities, the MexEF-OprN activity repressed RhlR activity. This is partially mediated by the delay in both PQS activation and the production of PqsE that would normally activate RhlR. There is an unidentified product regulated by MexEF-OprN that further represses RhlR activity. At higher cell densities, LasR activity counteracts the repression by MexEF-OprN and results in *rhlI/rhlR* and *pqsR* expression, leading to RhlR activity.

**Figure 6.**
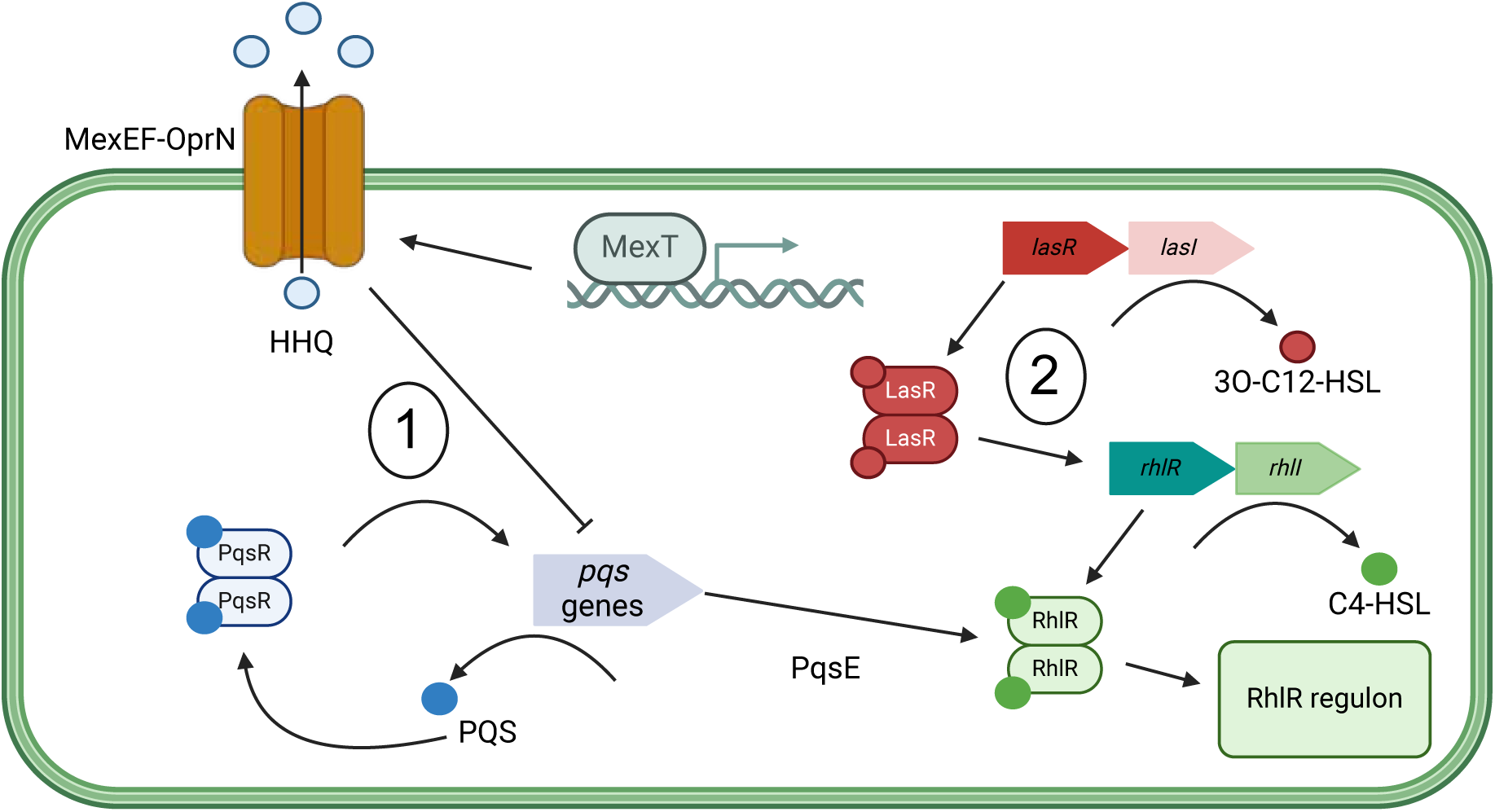
MexT negatively regulates RhlR activity, while LasR positively regulates *rhlR* expression. Cartoon showing the model of proposed MexT-mediated regulation of RhlR. At lower cell density, MexT positively regulates *mexEF-oprN*, delaying PQS QS and *pqsE* expression, and results in less active RhlR. At higher cell densities, LasR activity results in *rhlR* expression and an increase in RhlR activity.

### MexT is not the sole factor affecting QS architectures in P. aeruginosa

We wondered about the diversity of quorum sensing architectures and their regulation in other *P. aeruginosa* isolates. Recent work has identified clinical isolates that exhibit LasR-independent RhlR QS, but these had *lasR* null mutations at the times of isolation. It is possible that other mutations (such as in *mexT*) resulted in LasR-independent RhlR QS in these clinical isolates. We queried whether the LasR-dependent hierarchy was present in other isolates. We reasoned that if we knockout *lasR* in clinical isolates and RhlR activity is maintained, then these isolates encode a QS architecture different from PAO1. However, if we knockout *lasR* in clinical isolates and RhlR activity is abrogated, then the isolates display QS architecture akin to PAO1.

We started by assessing LasR and RhlR kinetics in three clinical isolates from chronic infections in people with cystic fibrosis (E192, E194, and E195) (17). These are isolates from the Early *Pseudomonas* Infection Control (EPIC) study (34) and harbor functional LasR proteins. Each isolate was transformed with the LasR and RhlR reporter plasmids described earlier and compared to wild-type PAO1 (**Figure 7**). All clinical isolates had active LasR QS that was similar to PAO1. In contrast, RhlR activity occurred earlier in all clinical isolates compared to PAO1. All three isolates exhibited at least 5-fold higher RhlR activity compared to PAO1. Together, these data show that the clinical isolates exhibit differences in QS compared to PAO1, possibly reflecting differences in QS architecture.

**Figure 7.**
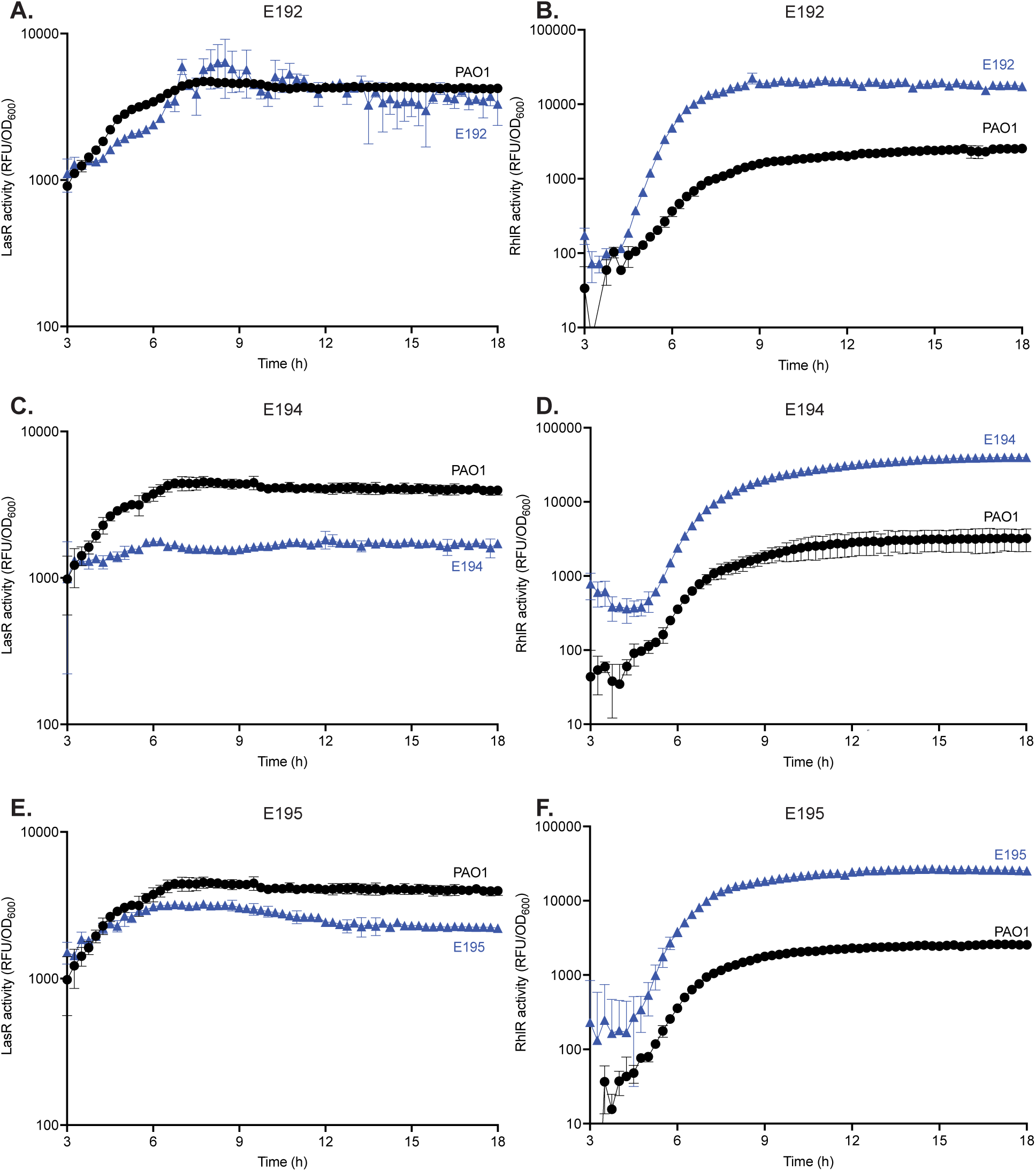
RhlR, but not LasR, activity is different between PAO1 and EPIC isolates. LasR activity for PAO1 compared to E192 (**A**), E194 (**C**), and E195 (**E**), measured using a LasR activity reporter plasmid. RhlR activity for PAO1 compared to E192 (**B**), E194 (**D**), and E195 (**F**), measured using a RhlR activity reporter plasmid.

We compared the QS architecture of clinical isolates to that of PAO1 by generating *lasR* deletion mutants and measuring RhlR activity (**Figure 8**). In all clinical isolates, RhlR activity occurred later and at a lower magnitude with *lasR* deletion compared to the parent (**Figure 8BCD**). However, clinical isolates still retained some RhlR activity, which differs from PAO1. Of the three isolates, E195Δ*lasR* retained the most RhlR activity. For isolates E192 and E194, RhlR activation in *lasR* mutants followed a linear pattern. For isolate E195, we observed RhlR activation in the *lasR* knockout followed an S-curve pattern similar to wild-type PAO1 and the parental strains.

**Figure 8.**
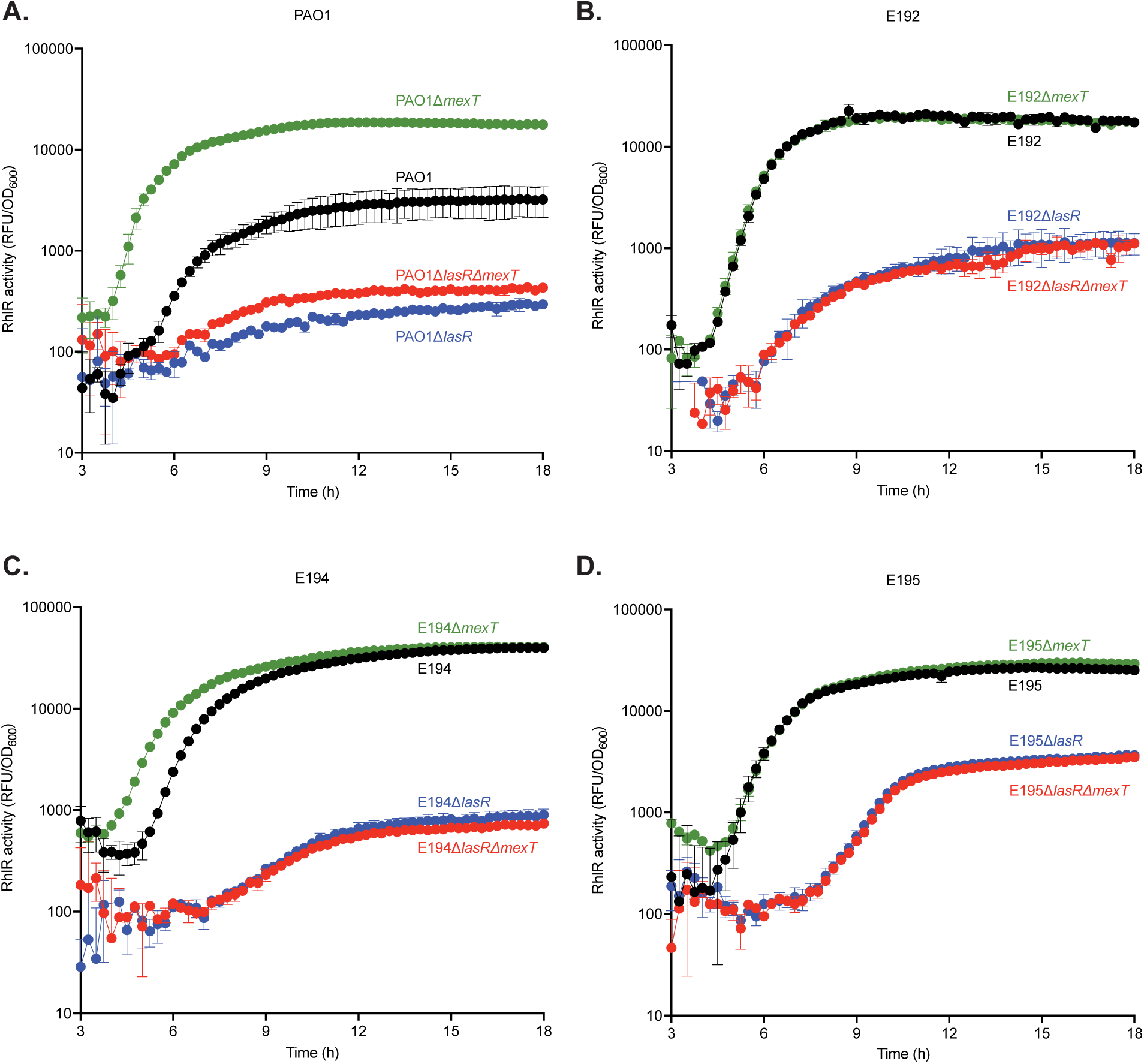
EPIC isolates exhibit LasR-dependent but MexT-independent RhlR activity. RhlR activity for wild-type, *lasR* and *mexT* knockouts, and *lasR* - *mexT* double knockout mutants for PAO1 (**A**), E192 (**B**), E194 (**C**), and E195 (**D**) measured using a RhlR activity reporter plasmid.

We next assessed whether the variability of LasR-dependence was due to MexT modulation of QS architecture. To ask this question, we performed whole genome sequencing on each clinical isolate to identify changes in the overall genome compared to PAO1. Each isolate encodes a larger genome than wild-type PAO1, ranging from 50 kbp to 600 kbp larger. E194 and E195 have identical *mexT* sequences to PAO1. E192 contained non-synonymous mutations that resulted in an I298F and D302E variant. We also assessed the MexT regulator, MexS, that is shown to be inactive in PAO1 due to a N249D mutation. In contrast to PAO1, all clinical isolates encode N249, indicating each has a functional MexS protein.

To investigate MexT activity in the clinical isolates, we transformed each isolate with a MexT reporter that contains the promoter of *mexE*, cloned upstream of *gfp* in an episomal plasmid. Compared to wild-type PAO1, MexT activity in each clinical isolate was almost undetectable (**Supplemental Figure 1**). MexT is activated due to oxidative stress, and others have demonstrated that the chemical compound diamide induces oxidative stress and activates MexT (22). PAO1 has constitutive MexT activity, but we observed a slight increase in MexT activity with the addition of diamide. In contrast, with the addition of diamide to each clinical isolate, we observed activation of MexT, indicating that MexT in each clinical isolate is functional, including the double variant encoded in E192.

We next determined if RhlR activity in the clinical isolates was modulated by MexT (as it is in PAO1) by generating *mexT* knockout mutants in each isolate and measuring RhlR activity (**Figure 8**). E192Δ*mexT* and E195Δ*mexT* displayed no differences in RhlR activity compared to wild-type isolates. Interestingly, E194Δ*mexT* showed earlier RhlR activity with the same terminal intensity compared to wild-type E194. Altogether, the clinical isolates differ from PAO1: the absence of MexT does not substantially change RhlR activity in *mexT* knockout mutants.

We asked whether MexT affects the relationship between LasR and RhlR QS. In PAO1, we observed an increase in RhlR QS in PAO1Δ*lasR*Δ*mexT* compared to PAO1Δ*lasR* (**Figure 8A**). In contrast, each clinical isolate exhibited no change in RhlR activity when comparing *lasR* knockout mutants to *lasR* and *mexT* double knockout mutants (**Figure 8BCD**). Together these results indicate that while the clinical isolates depend on LasR for RhlR activity, their RhlR activity is not negatively regulated by MexT as it is in PAO1.

## Discussion

In this work, we explored QS architectures in laboratory and clinical isolates of *P. aeruginosa* and how these architectures are modulated. We investigated how the transcription factor MexT promotes a LasR-dependent quorum sensing architecture in the laboratory strain PAO1 through bacterial genetics and transcriptomics. We discovered that the MexT-regulated genes *mexE*, *mexF*, and *oprN*, contribute to QS architecture. When these are inactivated, there is an increase in RhlR activity compared to the wild-type. Further, MexEF-OprN enforces this architecture through regulation of PQS QS and PqsE. By assessing the diversity of quorum sensing architecture across *P. aeruginosa*, we discovered that the QS hierarchy in a subset of clinical isolates from chronic infections of people with cystic fibrosis are LasR-dependent, but the degree of dependence varies across isolates, with none exhibiting absolute LasR dependence. Finally, we found that while there is LasR-dependent RhlR QS, in some isolates it was not dependent on MexT activity, demonstrating that the clinical isolates had a similar architecture but may depend on a modulator other than MexT.

Our study identifies the mechanism of a key regulator of QS in *P. aeruginosa*. We originally hypothesized that QS regulation was mediated either by the efflux pump MexEF-OprN or the chaperone PqsE. However, our data suggests it is the combination of both – induction of *mexEF-oprN* expression and repression of *pqsE* expression – that drives the LasR-dependent RhlR QS observed in strain PAO1. Several other studies have identified the QS signals C4-HSL, 3OC12-HSL, and HHQ as potential or established substrates of the efflux pump MexEF-OprN, supporting our finding that MexEF-OprN regulates QS (21, 30). Separate studies have shown that a suite of RhlR-regulated genes requires PqsE (32, 33). Our study shows a synergistic role between the two in regulating the *Pa* QS architecture, and we identified two potential mechanisms by which this happens. Further, our study reveals that RhlR-regulation by MexEF-OprN is also modulated by a PqsE-independent pathway. We are exploring potential candidates in a PqsE-independent pathway by examining the MexT regulon we identified in this study.

Our laboratory and others have characterized QS extensively in strain PAO1 (2, 3, 23, 35), but we wanted to further explore the diversity of QS architectures. We previously identified LasR-independent QS in clinical isolates derived from chronic infections of people with cystic fibrosis (15). However, these isolates harbored *lasR* null mutations at the time of isolation, and it was unknown if these isolates developed a different architecture from PAO1 due to other genomic differences. We showed in this study that clinical isolates can also have LasR-dependent QS, like PAO1. Surprisingly, we observed varying degrees of dependence on LasR, ranging from complete dependence on LasR for RhlR activity in E194 and partial dependence on LasR for RhlR activity in E195. It will be of interest to explore the genetic differences between the strains that result in differing degrees of dependence on LasR. Interestingly, while there was dependence on LasR for RhlR activity in all clinical isolates, we discovered that this was not dependent on MexT. This result illustrates that while PAO1 and clinical isolates share a similar quorum sensing architecture, the modulators seem to be different. Identifying these other modulators in the clinical isolates might illuminate the determinants of alternate QS architectures.

Our study of MexT-mediated regulation of QS affords us some insight into the formation of QS hierarchies. Some bacteria contain only a single QS circuit, while others have many (2, 36, 37). Our study begins to explore the purpose of having multiple QS circuits, through understanding how MexT functions to regulate QS. We also identified unique QS architectures in the EPIC isolates, indicating different regulators may be required for particular environments. MexT may have importance in the case of wound infections, where PAO1 was initially isolated, but it may be dispensable in the airways from which the EPIC isolates were obtained.

This work also offers insight into the plasticity of the interactions between QS circuits and how key regulators like MexT can alter QS architecture based on selective pressures and promote architectures that might be advantageous in specific environments. Further, by studying and identifying variants in regulators like MexT, we gain insight into how cooperative behaviors may function within and between individuals. It is easy to imagine that regulatory mutants can change QS architectures and cellular behaviors to allow for specialization within a community. For example, a *mexT* mutant may arise when RhlR activity is required early, while the wild-type defers the cost associated with RhlR QS, which might be advantageous when nutrients or other resources are limited.

Altogether, our study defined the regulation of a key modulator of *P. aeruginosa* growth and pathogenesis. By understanding how MexT modulates QS architecture, we begin to understand how bacterial sociality is controlled. These results have implications and applications that expand beyond bacterial pathogenesis and clinical settings. For example, how did MexT evolve to regulate quorum sensing architecture in PAO1, but not the clinical isolates? What selective factors favored genetic changes that resulted in LasR-dependent quorum sensing with different architecture regulators? Do the environment and other microbial interactions affect which architectures evolved (and which are beneficial in a given environment)? Our work pries the door open on understanding, where QS differs by interaction and environment, what the modulators of these QS architectures are.

## Materials and Methods

### Bacteria and growth conditions

Strains and plasmids used in this study are listed in **Supplemental Tables 3 and 4**. *P. aeruginosa* was grown in lysogeny broth (LB) buffered with 50 mM 3-(*N*-morpholino) propanesulfonic acid, pH 7.0 (LB-MOPS). *Escherichia coli* was grown in LB. Cultures were grown in 18-mm test tubes at a volume of 3 mL in a shaking incubator (250 RPM) at 37°C. For individual colony growth we used LB supplemented with 1.5% agar. Where required, broth cultures of *E. coli* and *P. aeruginosa* were supplemented with gentamicin at a concentration of 10 µg per mL (Gm10). *E. coli* colonies were grown on LB supplemented with 1.5% agar and gentamicin at 10 µg per mL. *P. aeruginosa* colonies were grown on LB supplemented with 1.5% agar and gentamicin at 100 µg per mL (Gm100) for transformations or 10 µg per mL for maintaining cultures.

### Construction of *P. aeruginosa* mutants

In all experiments with the laboratory strain PAO1, we used strain *P. aeruginosa* PAO1-UW (38). Clinical *P. aeruginosa* isolates were obtained from the EPIC Observational Study and are from children with cystic fibrosis 5-12 years of age (34). In-frame deletions of *lasR*, *pqsE*, *mexT*, and *mexEF* were generated using two-step allelic exchange as previously described (39). Briefly, constructs for gene deletions were created by using a pEXG2 vector backbone and Gibson assembly to 1000 bp of DNA flanking each side of the gene of interest to facilitate homologous recombination. *E. coli* S17-1 was transformed with each construct and used to deliver knockout plasmids to *P. aeruginosa* via conjugation. Merodiploids were selected by plating on *Pseudomonas* Isolate agar containing Gm100, and deletion mutants were then selected on LB agar containing 10-15% sucrose and no sodium chloride. All deletion mutants were confirmed by PCR and sequencing of genomic DNA.

Overexpression constructs were created using a pUC18T mini Tn7T integrating plasmid with an arabinose-inducible promoter and Gibson assembly to clone in either *pqsE* or *mexEF-oprN* CDS downstream of the promoter (40). The overexpression construct was electrotransformed into *P. aeruginosa* as described elsewhere and selected using LB supplemented with 1.5% agar and gentamicin at 100 µg per mL (Gm100) (41). Proper integration at the attTn7 attachment site was confirmed by PCR and sequencing of genomic DNA. Gentamicin susceptibility was restored using Flp-mediated excision of gentamicin resistance marker (42).

Reporter plasmids pP*_lasI_*-*gfp*, pP*_rhlA_*-*gfp*, pP*_pqsA_*-*gfp,* and pP*_mexE_*-*gfp* were constructed in pBBR1MCS-5 and contain about 200-500 bp upstream of each gene fused to *gfp*. Plasmids were electrotransformed into *P. aeruginosa* as described above.

### Assays for LasR, RhlR, and PqsR activity

To measure LasR activity, we inoculated individual duplicate colonies of strains with the transcriptional reporter P*_lasI_*-*gfp* into LB-MOPS supplemented with Gm10 and incubated at 37°C with shaking for 18 hours. Cultures were back diluted to an OD_600_ of 0.01 and grown until OD_600_ of 0.1. Exponential phase cultures were used to inoculate a 48-well plate to an OD_600_ of 0.01 with a 300 µL final volume. Plates were incubated at 37°C with shaking for 18 hours in a Biotek Synergy H1 microplate reader. GFP fluorescence (excitation 485 nm, emission 535 nm) and OD_600_ were measured every 20 minutes. The plasmids pP*_rhlA_*-*gfp* and pP*_pqsA_*-*gfp* were similarly introduced into each strain to measure RhlR and PqsR activity, respectively. Where appropriate, 20 µM final PQS (in methanol) was added before plates were incubated in the microplate reader. Similarly, to induce *pqsE* and *mexEF-oprN* expression, LB-MOPS supplemented with Gm10 was inoculated with a final concentration of 0.1% arabinose before plates were incubated in the microplate reader. All experiments were performed in triplicate.

### Assay for MexT activity

Strains were transformed with the pP*_mexE_*-*gfp* reporter plasmid (22). Single colonies were used to inoculate LB-MOPS supplemented with Gm10 and grown for 20 hours. Cultures were back diluted to an OD_600_ of 0.05 in LB-MOPS supplemented with Gm10. To determine whether MexT activity can be activated, 8 mM final diamide [1,1’-Azobis(N,N-dimethylformamide), TCI America] was added to cultures. Cultures were incubated for 18 hours at 37°C with shaking. Endpoint GFP fluorescence (excitation 485 nm, emission 535 nm) and OD_600_ were measured using a Biotek Synergy H1 microplate reader. For all endpoint measurements, the background fluorescence of LB was subtracted from the final fluorescence value of the reporter culture.

### RNA isolation and RNA sequencing

Wild-type PAO1 and PAO1Δ*mexT* cultures were started from single colonies in LB-MOPS in biological triplicate. Cultures were incubated for 18 hours at 37°C with shaking. Cultures were back diluted to an OD_600_ of 0.1 until cultures doubled, and then cultures are diluted back to an OD_600_ of 0.05. At an OD_600_ of 1.0 and 2.0, a total of an OD_600_ of 2.0 was pelleted at 4000 RPM for 5 min. Supernatant was discarded and pellets were resuspended in 1 mL QIAzol containing lysis beads. Samples were lysed by bead-beating at maximum RPM for 1 min with chilling for 5 min. Bead-beating was repeated twice. Chloroform was added, shaken vigorously, and centrifuged for 15 min at 12,000 x g at 4°C. 450 µL of the upper phase was combined with 675 µL of 100% ethanol. Samples RNA was extracted using a RNeasy kit with one on-column DNase (cat. No. 79254, Qiagen) treatment, and RNA was eluted using RNase-free water.

rRNA depletion, library generation, and >20 million 150-bp paired-end Illumina HiSeq reads were generated for each sample commercially (Genewiz; Azenta). Trimmomatic with default settings (https://github.com/timflutre/trimmomatic) was used to trim adapters prior to alignment against the PAO1 reference genome (accession NC_002516) using StrandNGS version 3.3.1 (Strand Life Sciences, Bangalore, India). DESeq2 (27) was used for differential expression analysis, using the Benjamini-Hochberg adjustment for multiple comparisons and a false-discovery rate α = 0.05. Samples were grouped according to each strain and growth phase for the DESeq2 differential expression analyses. MexT regulons were determined by comparing the wild-type PAO1 to PAO1Δ*mexT* for a given growth phase and imposing a 2-fold minimal fold change threshold. RhlR regulon genes were filtered out based on a compiled RhlR core regulon (15, 16, 28). The RNA-seq data are deposited in the Gene Expression Omnibus under accession number GSE291609.

### qRT PCR analysis

RNA was extracted as described above. 200 ng total cDNA was generated by reverse transcription using the qScript cDNA synthesis kit. Quantitative real-time PCR was performed using 2.5 ng total cDNA using the PowerTrack SYBR Green Mix. Primers targeting *mexE* were used to monitor MexT activity. Primers amplifying *rplU* were used as a housekeeping gene. Data analysis was determined using relative gene expression levels using 2^-ΔΔCt^ method.

### Measurement of C4-HSL concentrations

Individual colonies of wild-type PAO1 or PAO1Δ*mexEF* were inoculated into 3 mL LB-MOPS in triplicate. Cultures were grown at 37°C in a shaking incubator for 18 hours. Cultures were back diluted to an OD_600_ of 0.1 and grown until OD_600_ of 0.2, and then back diluted to OD_600_ of 0.1. At OD_600_ of 0.3, 0.8, and 2.0, cultures were removed to extract C4-HSL. To extract total C4-HSL, 3 mL of culture was transferred to an 18-mm glass test tube and mixed with 3 mL acidified ethyl acetate. Samples were vortexed for 30 seconds and rested for 10 minutes to allow for phase separation. The clear top layer was extracted to a 10 mL glass collection tube. The extraction was repeated and added to the same 10 mL glass collection tube. To extract extracellular C4-HSL, we first centrifuged 3 mL of culture at 5000xg for 5 min and collected the supernatant in an 18 mm glass test tube and mixed with 3 mL acidified ethyl acetate. Samples were vortexed on high for 30 seconds and rested for 10 minutes to allow for phase separation. The clear top layer was extracted to a 10 mL glass collection tube. The extraction was repeated and added to the same 10 mL glass collection tube. For each sample, acidified ethyl acetate was evaporated using an N_2_ evaporator until no liquid remained. Dried fractions were resuspended in 300 µL ethyl acetate to create a 10x stock.

To measure C4-HSL, we used a bioassay strain E. coli DH5α/pEDP61.5 containing *tacp rhlR* and a *rhlAB-lacZ* reporter (43). The bioassay strain was grown in 5 mL LB supplemented with 100 µg per mL carbenicillin for 18 hours at 37°C with shaking. Cultures were back diluted to OD_600_ of 0.05 and grown until an OD_600_ of 0.1-0.2. Cultures were induced with 1 mM final IPTG and grown to OD_600_ of 0.5. C4-HSL standards and the above extracted samples were deposited into 1.5 mL microcentrifuge tubes and dried with N_2_. When cultures reach an OD_600_ of 0.5, 0.5 mL of culture was added to the prepared signal-containing tubes. Cultures were grown for 3 hours at 37°C with shaking. Cultures were lysed with the addition of 50 µL chloroform, vortexed, and 10 µL of the top phase of each sample was transferred to a 96-well white microtiter Optiplate. Luminescence was measured using the Galacton-Plus chemiluminescent kit.

## Acknowledgements

This research was supported by NIH grant R01 AI177575 (to A.A.D.). A.F. was supported by a Cystic Fibrosis Foundation Postdoctoral Fellowship (FRANDO24F0) and an NIH/NIAID training grant (T32 HD007233-42).

## References

1. Miller MB, Bassler BL. 2001. Quorum sensing in bacteria. Annu Rev Microbiol 55:165–99.

2. Schuster M, Greenberg EP. 2006. A network of networks: quorum-sensing gene regulation in *Pseudomonas aeruginosa*. Int J Med Microbiol 296:73–81.

3. Schuster M, Lostroh CP, Ogi T, Greenberg EP. 2003. *Identification, timin*g, and signal specificity of Pseudomonas aeruginosa quorum-controlled genes: a transcriptome analysis. J Bacteriol 185:2066–79.

4. Fuqua WC, Winans SC, Greenberg EP. 1994. Quorum sensing in bacteria: the LuxR-LuxI family of cell density-responsive transcriptional regulators. J Bacteriol 176:269–75.

5. Deziel E, Lepine F, Milot S, He J, Mindrinos MN, Tompkins RG, Rahme LG. 2004. Analysis of *Pseudomonas aeruginosa* 4-hydroxy-2-alkylquinolines (HAQs) reveals a role for 4-hydroxy-2-heptylquinoline in cell-to-cell communication. Proc Natl Acad Sci U S A 101:1339–44.

6. Gallagher LA, McKnight SL, Kuznetsova MS, Pesci EC, Manoil C. 2002. Functions required for extracellular quinolone signaling by *Pseudomonas aeruginosa*. J Bacteriol 184:6472–80.

7. Pesci EC, Milbank JB, Pearson JP, McKnight S, Kende AS, Greenberg EP, Iglewski BH. 1999. Quinolone signaling in the cell-to-cell communication system of *Pseudomonas aeruginosa*. Proc Natl Acad Sci U S A 96:11229–34.

8. Diggle SP, Cornelis P, Williams P, Camara M. 2006. 4-quinolone signalling in Pseudomonas aeruginosa: old molecules, new perspectives. Int J Med Microbiol 296:83–91.

9. Scholz RL, Greenberg EP. 2017. Positive autoregulation of an acyl-homoserine lactone quorum-sensing circuit synchronizes the population response. mBio 8.

10. de Kievit TR, Kakai Y, Register JK, Pesci EC, Iglewski BH. 2002. Role of the *Pseudomonas aeruginosa las* and *rhl* quorum-sensing systems in *rhlI* regulation. FEMS Microbiol Lett 212:101–6.

11. Sandoz KM, Mitzimberg SM, Schuster M. 2007. Social cheating in *Pseudomonas aeruginosa* quorum sensing. Proc Natl Acad Sci U S A 104:15876–81.

12. Borgert SR, Henke S, Witzgall F, Schmelz S, Zur Lage S, Hotop SK, Stephen S, Lubken D, Kruger J, Gomez NO, van Ham M, Jansch L, Kalesse M, Pich A, Bronstrup M, Haussler S, Blankenfeldt W. 2022. Moonlighting chaperone activity of the enzyme PqsE contributes to RhlR-controlled virulence of *Pseudomonas aeruginosa*. Nat Commun 13:7402.

13. Feathers JR, Richael EK, Simanek KA, Fromme JC, Paczkowski JE. 2022. Structure of the RhlR-PqsE complex from Pseudomonas aeruginosa reveals mechanistic insights into quorum-sensing gene regulation. Structure 30:1626–1636 e4.

14. Brouwer S, Pustelny C, Ritter C, Klinkert B, Narberhaus F, Haussler S. 2014. The PqsR and RhlR transcriptional regulators determine the level of *Pseudomonas* quinolone signal synthesis in *Pseudomonas aeruginosa* by producing two different *pqsABCDE* mRNA isoforms. J Bacteriol 196:4163–71.

15. Asfahl KL, Smalley NE, Chang AP, Dandekar AA. 2022. Genetic and transcriptomic characteristics of RhlR-dependent quorum sensing in cystic fibrosis isolates of *Pseudomonas aeruginosa*. mSystems 7:e0011322.

16. Cruz RL, Asfahl KL, Van den Bossche S, Coenye T, Crabbe A, Dandekar AA. 2020. RhlR-regulated acyl-homoserine lactone quorum sensing in a cystic fibrosis isolate of *Pseudomonas aeruginosa*. mBio 11.

17. Feltner JB, Wolter DJ, Pope CE, Groleau MC, Smalley NE, Greenberg EP, Mayer-Hamblett N, Burns J, Deziel E, Hoffman LR, Dandekar AA. 2016. LasR variant cystic fibrosis isolates reveal an adaptable quorum-sensing hierarchy in *Pseudomonas aeruginosa*. mBio 7.

18. Sobel ML, Neshat S, Poole K. 2005. Mutations in PA2491 (*mexS*) promote MexT-dependent mexEF-oprN expression and multidrug resistance in a clinical strain of *Pseudomonas aeruginosa*. J Bacteriol 187:1246–53.

19. Lee S, Gallagher L, Manoil C. 2021. Reconstructing a wild-type *Pseudomonas aeruginosa* reference strain PAO1. J Bacteriol 203:e0017921.

20. Richardot C, Juarez P, Jeannot K, Patry I, Plesiat P, Llanes C. 2016. Amino acid substitutions account for most MexS alterations in clinical *nfxC* mutants of *Pseudomonas aeruginosa*. Antimicrob Agents Chemother 60:2302–10.

21. Lamarche MG, Deziel E. 2011. MexEF-OprN efflux pump exports the *Pseudomonas* quinolone signal (PQS) precursor HHQ (4-hydroxy-2-heptylquinoline). PLoS One 6:e24310.

22. Kostylev M, Smalley NE, Chao MH, Greenberg EP. 2023. Relationship of the transcription factor MexT to quorum sensing and virulence in *Pseudomonas aeruginosa*. J Bacteriol 205:e0022623.

23. Kostylev M, Kim DY, Smalley NE, Salukhe I, Greenberg EP, Dandekar AA. 2019. Evolution of the *Pseudomonas aeruginosa* quorum-sensing hierarchy. Proc Natl Acad Sci U S A 116:7027–7032.

24. Oshri RD, Zrihen KS, Shner I, Omer Bendori S, Eldar A. 2018. Selection for increased quorum-sensing cooperation in *Pseudomonas aeruginosa* through the shut-down of a drug resistance pump. ISME J 12:2458–2469.

25. Tian ZX, Fargier E, Mac Aogain M, Adams C, Wang YP, O’Gara F. 2009. Transcriptome profiling defines a novel regulon modulated by the LysR-type transcriptional regulator MexT in *Pseudomonas aeruginosa*. Nucleic Acids Res 37:7546–59.

26. Kohler T, Epp SF, Curty LK, Pechere JC. 1999. Characterization of MexT, the regulator of the MexE-MexF-OprN multidrug efflux system of *Pseudomonas aeruginosa*. J Bacteriol 181:6300–5.

27. Love MI, Huber W, Anders S. 2014. Moderated estimation of fold change and dispersion for RNA-seq data with DESeq2. Genome Biol 15:550.

28. Chugani S, Kim BS, Phattarasukol S, Brittnacher MJ, Choi SH, Harwood CS, Greenberg EP. 2012. Strain-dependent diversity in the *Pseudomonas aeruginosa* quorum-sensing regulon. Proc Natl Acad Sci U S A 109:E2823–31.

29. Kohler T, Michea-Hamzehpour M, Henze U, Gotoh N, Curty LK, Pechere JC. 1997. Characterization of MexE-MexF-OprN, a positively regulated multidrug efflux system of *Pseudomonas aeruginosa*. Mol Microbiol 23:345–54.

30. Kristensen R, Andersen JB, Rybtke M, Jansen CU, Fritz BG, Kiilerich RO, Uhd J, Bjarnsholt T, Qvortrup K, Tolker-Nielsen T, Givskov M, Jakobsen TH. 2024. Inhibition of *Pseudomonas aeruginosa* quorum sensing by chemical induction of the MexEF-oprN efflux pump. Antimicrob Agents Chemother 68:e0138723.

31. Yamaguchi A, Nakashima R, Sakurai K. 2015. Structural basis of RND-type multidrug exporters. Front Microbiol 6:327.

32. Simanek KA, Taylor IR, Richael EK, Lasek-Nesselquist E, Bassler BL, Paczkowski JE. 2022. The PqsE-RhlR interaction regulates RhlR DNA Binding to control virulence factor production in *Pseudomonas aeruginosa*. Microbiol Spectr 10:e0210821.

33. Letizia M, Mellini M, Fortuna A, Visca P, Imperi F, Leoni L, Rampioni G. 2022. PqsE expands and differentially modulates the RhlR quorum sensing regulon in *Pseudomonas aeruginosa*. Microbiol Spectr 10:e0096122.

34. Rosenfeld M, Emerson J, McNamara S, Joubran K, Retsch-Bogart G, Graff GR, Gutierrez HH, Kanga JF, Lahiri T, Noyes B, Ramsey B, Ren CL, Schechter M, Morgan W, Gibson RL, Sites ESGPC. 2010. Baseline characteristics and factors associated with nutritional and pulmonary status at enrollment in the cystic fibrosis EPIC observational cohort. Pediatr Pulmonol 45:934–44.

35. Miranda SW, Asfahl KL, Dandekar AA, Greenberg EP. 2022. *Pseudomonas aeruginosa* quorum sensing. Adv Exp Med Biol 1386:95–115.

36. Engebrecht J, Nealson K, Silverman M. 1983. Bacterial bioluminescence: isolation and genetic analysis of functions from *Vibrio fischeri*. Cell 32:773–81.

37. Le Guillouzer S, Groleau MC, Deziel E. 2017. The Complex Quorum Sensing Circuitry of Burkholderia thailandensis Is Both Hierarchically and Homeostatically Organized. mBio 8.

38. Stover CK, Pham XQ, Erwin AL, Mizoguchi SD, Warrener P, Hickey MJ, Brinkman FS, Hufnagle WO, Kowalik DJ, Lagrou M, Garber RL, Goltry L, Tolentino E, Westbrock-Wadman S, Yuan Y, Brody LL, Coulter SN, Folger KR, Kas A, Larbig K, Lim R, Smith K, Spencer D, Wong GK, Wu Z, Paulsen IT, Reizer J, Saier MH, Hancock RE, Lory S, Olson MV. 2000. Complete genome sequence of *Pseudomonas aeruginosa* PAO1, an opportunistic pathogen. Nature 406:959–64.

39. Hmelo LR, Borlee BR, Almblad H, Love ME, Randall TE, Tseng BS, Lin C, Irie Y, Storek KM, Yang JJ, Siehnel RJ, Howell PL, Singh PK, Tolker-Nielsen T, Parsek MR, Schweizer HP, Harrison JJ. 2015. Precision-engineering the *Pseudomonas aeruginosa* genome with two-step allelic exchange. Nat Protoc 10:1820–41.

40. Choi KH, Schweizer HP. 2006. mini-Tn7 insertion in bacteria with single attTn7 sites: example *Pseudomonas aeruginosa*. Nat Protoc 1:153–61.

41. Choi KH, Kumar A, Schweizer HP. 2006. A 10-min method for preparation of highly electrocompetent *Pseudomonas aeruginosa* cells: application for DNA fragment transfer between chromosomes and plasmid transformation. J Microbiol Methods 64:391–7.

42. Hoang TT, Karkhoff-Schweizer RR, Kutchma AJ, Schweizer HP. 1998. A broad-host-range Flp-FRT recombination system for site-specific excision of chromosomally-located DNA sequences: application for isolation of unmarked *Pseudomonas aeruginosa* mutants. Gene 212:77–86.

43. Pearson JP, Pesci EC, Iglewski BH. 1997. Roles of *Pseudomonas aeruginosa las* and *rhl* quorum-sensing systems in control of elastase and rhamnolipid biosynthesis genes. J Bacteriol 179:5756–67.

