## Supplemental Figure 1 for "Modulation of the *Pseudomonas aeruginosa* quorum sensing cascade by MexT-regulated factors"

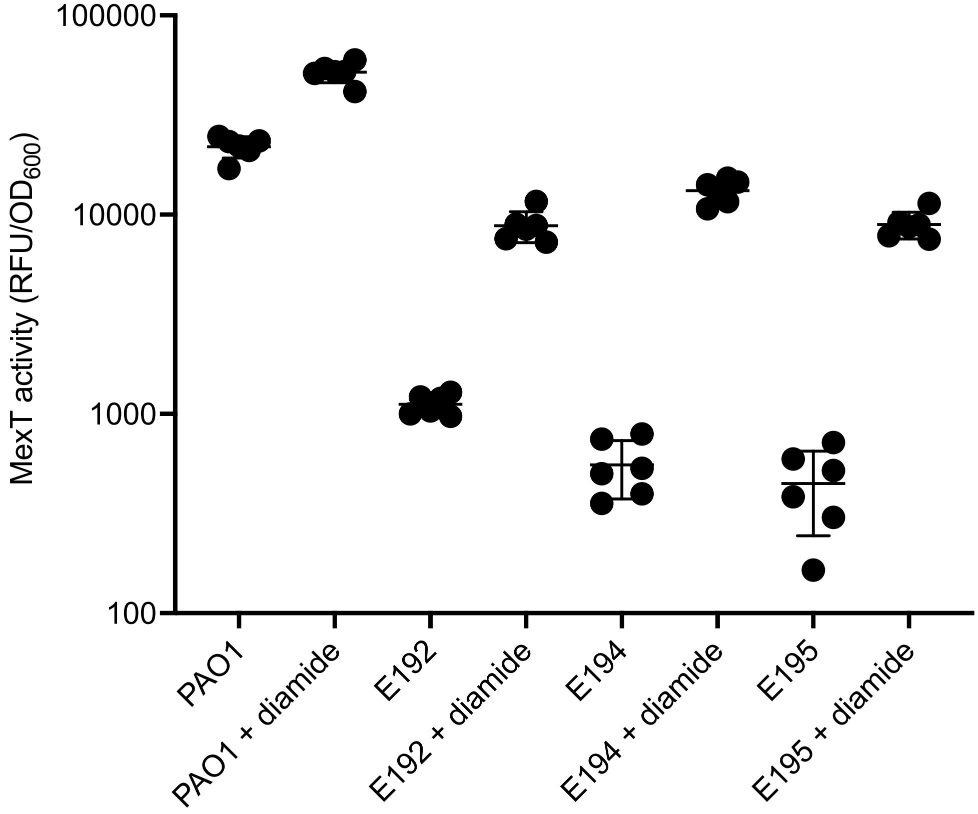


**Supplemental Figure 1**. **EPIC isolates have functional MexT that can be activated with oxidative stress**. MexT activity was measured PAO1, E192, E194, and E195 with and without the addition of 8 mM diamide using a MexT activity reporter plasmid. MexT activity was measured after 18 hours. The y-axis shows RFU normalized to $OD_{600}$.
