## Supplemental Table 1 for "Modulation of the *Pseudomonas aeruginosa* quorum sensing cascade by MexT-regulated factors"

**Supplemental Table 1.** MexT regulon at an $OD_{600}$ of 1.0.

| Gene Symbol | Locus Tag | Gene ID | log2FC_MexT_OD1_WT_OD1 | pVal_MexT_OD1_WT_OD1 | pAdj_MexT_OD1_WT_OD1 |
|---|---|---|---|---|---|
| bkdA1 | PA2247 | 877901 | 1.0935633 | 0.000626119 | 0.014965046 |
| bkdA2 | PA2248 | 879226 | 1.2740743 | 7.6277E-12 | 5.89982E-10 |
| chiC | PA2300 | 878587 | 1.0064931 | 3.53927E-05 | 0.001187363 |
| fabH2 | PA3333 | 882498 | 4.3242774 | 2.521E-42 | 5.39957E-40 |
| hcnA | PA2193 | 882197 | 1.9333028 | 4.78457E-17 | 4.67461E-15 |
| hcnB | PA2194 | 882182 | 2.147197 | 6.7129E-33 | 1.09953E-30 |
| hcnC | PA2195 | 882196 | 2.0661895 | 3.06163E-16 | 2.79512E-14 |
| hcpA | PA1512 | 883081 | 1.5819535 | 1.12874E-07 | 5.93016E-06 |
| hcpB | PA5267 | 879579 | 1.6977073 | 1.04014E-07 | 5.56973E-06 |
| hcpC | PA0263 | 880866 | 1.223545 | 4.33761E-05 | 0.00142095 |
| hisJ | PA2923 | 882695 | 1.7895652 | 0.000665066 | 0.015507184 |
| hpcG | PA4127 | 880260 | 1.2797428 | 1.84235E-14 | 1.57846E-12 |
| kdpB | PA1634 | 883011 | 1.342634 | 2.83956E-07 | 1.43564E-05 |
| kdpC | PA1635 | 883003 | 1.1208234 | 0.001491754 | 0.030883197 |
| ladS | PA3974 | 878890 | 1.3900806 | 3.81084E-28 | 5.17623E-26 |
| lasB | PA3724 | 880368 | 1.7282188 | 1.88026E-16 | 1.77477E-14 |
| lecB | PA3361 | 882528 | 2.262034 | 2.69701E-24 | 3.26514E-22 |
| mexH | PA4206 | 880450 | 1.3874915 | 5.90182E-09 | 3.65192E-07 |
| opdC | PA0162 | 879518 | 1.328185 | 4.22191E-18 | 4.27488E-16 |
| opmD | PA4208 | 880347 | 1.3372176 | 3.44409E-06 | 0.000146413 |
| oprB | PA3186 | 882788 | 1.0349458 | 2.51372E-10 | 1.77201E-08 |
| oprD | PA0958 | 881970 | 1.9223132 | 1.88749E-29 | 2.62785E-27 |
| PA0050 | PA0050 | 878575 | 1.2353244 | 1.46733E-06 | 6.64356E-05 |
| PA0122 | PA0122 | 880611 | 1.9986491 | 1.4E-45 | 3E-43 |
| PA0165 | PA0165 | 879418 | 2.2565334 | 4.682E-35 | 7.90123E-33 |
| PA1131 | PA1131 | 878079 | 2.1131742 | 1.01013E-26 | 1.25009E-24 |
| PA1221 | PA1221 | 878373 | 1.5249729 | 2.78258E-06 | 0.000120125 |
| PA1657 | PA1657 | 879559 | 1.1738352 | 0.000180967 | 0.005064359 |
| PA1658 | PA1658 | 879592 | 1.0103709 | 0.001734925 | 0.033664793 |
| PA1668 | PA1668 | 882100 | 1.4235904 | 1.48754E-08 | 8.81291E-07 |
| PA1669 | PA1669 | 882077 | 1.4195819 | 6.22292E-13 | 4.88105E-11 |
| PA1869 | PA1869 | 881955 | 2.6896315 | 1.40537E-38 | 2.69879E-36 |
| PA2067 | PA2067 | 878262 | 1.5559263 | 0.000102977 | 0.003116728 |
| PA2068 | PA2068 | 877702 | 1.500314 | 2.86148E-07 | 1.43564E-05 |
| PA2069 | PA2069 | 877951 | 1.8476362 | 1.46466E-13 | 1.18213E-11 |
| PA2097 | PA2097 | 877933 | 1.0474402 | 0.000499588 | 0.012149375 |
| PA2260 | PA2260 | 877758 | 1.7568915 | 2.3178E-09 | 1.49348E-07 |

| | | | | | |
|---|---|---|---|---|---|
| PA2322 | PA2322 | 883019 | 1.2440337 | 3.1382E-05 | 0.00109229 |
| PA2330 | PA2330 | 883026 | 1.063155 | 2.43446E-08 | 1.39768E-06 |
| PA2362 | PA2362 | 882222 | 1.054405 | 0.001591969 | 0.032007545 |
| PA2781 | PA2781 | 882770 | 1.5935738 | 5.93892E-10 | 4.08319E-08 |
| PA2782 | PA2782 | 882771 | 1.4813118 | 1.04252E-06 | 4.8788E-05 |
| PA3325 | PA3325 | 882490 | 1.9513874 | 1.0339E-27 | 1.3709E-25 |
| PA3326 | PA3326 | 882491 | 2.6452918 | 8.64049E-30 | 1.23382E-27 |
| PA3327 | PA3327 | 882492 | 4.108884 | 0 | 0 |
| PA3328 | PA3328 | 882493 | 5.0676017 | 0 | 0 |
| PA3329 | PA3329 | 882494 | 4.3763924 | 0 | 0 |
| PA3330 | PA3330 | 882495 | 4.6232724 | 0 | 0 |
| PA3331 | PA3331 | 882496 | 3.8526144 | 6.94827E-38 | 1.24822E-35 |
| PA3332 | PA3332 | 882497 | 4.808917 | 0 | 1.63E-43 |
| PA3334 | PA3334 | 882499 | 4.9308286 | 1.2638E-39 | 2.5136E-37 |
| PA3335 | PA3335 | 882500 | 3.0825014 | 3.38027E-24 | 4.00526E-22 |
| PA3377 | PA3377 | 877807 | 1.0674754 | 8.79978E-05 | 0.002707512 |
| PA3519 | PA3519 | 878876 | 1.8099405 | 0.000279836 | 0.007492329 |
| PA3574a | PA3574a | 17373374 | 1.5329958 | 0.000178185 | 0.005037107 |
| PA3677 | PA3677 | 879603 | 1.130277 | 3.5222E-05 | 0.001187363 |
| PA4128 | PA4128 | 881843 | 1.8722057 | 2.42235E-21 | 2.59424E-19 |
| PA4129 | PA4129 | 881844 | 1.8670844 | 1.93197E-23 | 2.19574E-21 |
| PA4130 | PA4130 | 881847 | 1.6660838 | 5.1973E-41 | 1.072E-38 |
| PA4131 | PA4131 | 881848 | 1.6437137 | 5.09822E-38 | 9.46399E-36 |
| PA4132 | PA4132 | 880050 | 1.2982695 | 1.20078E-13 | 9.83402E-12 |
| PA4133 | PA4133 | 880051 | 2.6578617 | 7.62808E-30 | 1.11792E-27 |
| PA4134 | PA4134 | 880084 | 1.532603 | 2.7568E-13 | 2.19323E-11 |
| PA4141 | PA4141 | 880096 | 2.1929696 | 8.603E-27 | 1.11419E-24 |
| PA4142 | PA4142 | 880112 | 1.696943 | 3.45337E-16 | 3.10191E-14 |
| PA4218 | PA4218 | 880532 | 1.3060786 | 4.56003E-07 | 2.22761E-05 |
| PA4219 | PA4219 | 880533 | 1.1179392 | 2.8783E-05 | 0.001014511 |
| PA4341 | PA4341 | 881469 | 1.0708456 | 0.001633992 | 0.03220807 |
| PA4596 | PA4596 | 881060 | 1.8199911 | 0.000672953 | 0.015615305 |
| PA5180 | PA5180 | 881723 | 1.2841951 | 3.50861E-05 | 0.001187363 |
| PA5220 | PA5220 | 882292 | 2.158614 | 1.40899E-19 | 1.48051E-17 |
| PA5328 | PA5328 | 878251 | 2.1779 | 0.000694229 | 0.015975863 |
| pchA | PA4231 | 881821 | 1.0329945 | 2.45362E-05 | 0.000881561 |
| pchB | PA4230 | 881846 | 1.7991292 | 2.26172E-09 | 1.48183E-07 |
| pchC | PA4229 | 881845 | 1.7834198 | 2.33314E-09 | 1.49348E-07 |
| pchD | PA4228 | 880566 | 1.197166 | 1.2456E-05 | 0.000478395 |

| | | | | | |
|---|---|---|---|---|---|
| pchE | PA4226 | 880048 | 1.4325575 | 4.90574E-14 | 4.1088E-12 |
| pchF | PA4225 | 880083 | 1.7178838 | 1.40273E-05 | 0.000527824 |
| pcrG | PA1705 | 879383 | 2.187794 | 2.97974E-06 | 0.000127647 |
| phnA | PA1001 | 878421 | 1.4487439 | 9.72031E-12 | 7.4154E-10 |
| phzA1 | PA4210 | 880515 | 3.0362558 | 1.94145E-08 | 1.12624E-06 |
| phzA2 | PA1899 | 878477 | 2.2792864 | 5.00004E-11 | 3.66385E-09 |
| phzB1 | PA4211 | 880518 | 3.5803676 | 0 | 0 |
| phzB2 | PA1900 | 878211 | 1.6081278 | 5.44639E-07 | 2.63747E-05 |
| phzC1 | PA4212 | 880519 | 3.7776842 | 3.57356E-06 | 0.000150766 |
| phzD2 | PA1902 | 881873 | 1.4399945 | 1.42108E-05 | 0.000531141 |
| phzE2 | PA1903 | 882029 | 1.0900362 | 7.57924E-06 | 0.000310359 |
| phzF2 | PA1904 | 879183 | 1.200616 | 4.80701E-05 | 0.001565512 |
| phzS | PA4217 | 881836 | 1.0434259 | 1.59601E-05 | 0.000592545 |
| popN | PA1698 | 879266 | 1.2545154 | 9.0131E-06 | 0.000355986 |
| pqsA | PA0996 | 880760 | 1.3597804 | 3.65007E-17 | 3.62986E-15 |
| pqsB | PA0997 | 883098 | 1.3997477 | 2.58712E-09 | 1.63724E-07 |
| pqsC | PA0998 | 880660 | 1.3641305 | 2.32949E-07 | 1.19138E-05 |
| pqsD | PA0999 | 880625 | 1.2042456 | 2.40565E-06 | 0.000104665 |
| pqsE | PA1000 | 880721 | 1.2848428 | 2.33184E-07 | 1.19138E-05 |
| pscE | PA1718 | 881908 | 1.1178021 | 0.000263567 | 0.007134711 |
| pscF | PA1719 | 879634 | 1.4003649 | 2.98472E-05 | 0.001045404 |
| qteE | PA2593 | 882299 | 1.2345799 | 1.98402E-11 | 1.4732E-09 |
| rbsA | PA1947 | 880680 | 1.6131611 | 3.62701E-05 | 0.00120951 |
| rbsC | PA1948 | 880795 | 1.4636799 | 5.64542E-08 | 3.18763E-06 |
| rhlA | PA3479 | 878955 | 2.9713337 | 0 | 0 |
| rhlI | PA3476 | 878967 | 1.3450693 | 6.14641E-16 | 5.43323E-14 |
| stk1 | PA1671 | 883070 | 1.1569456 | 1.85967E-08 | 1.09016E-06 |
| stp1 | PA1670 | 882277 | 1.3612847 | 5.66664E-08 | 3.18763E-06 |
| bexR | PA2432 | 882852 | -1.1018038 | 4.94325E-14 | 4.1088E-12 |
| gbuA | PA1421 | 881747 | -2.6100378 | 2.22422E-16 | 2.06445E-14 |
| mexE | PA2493 | 880212 | -6.275341 | 0 | 0 |
| mexF | PA2494 | 882884 | -6.372459 | 0 | 0 |
| mexT | PA2492 | 880417 | -6.5968676 | 0 | 6.7E-44 |
| moaA1 | PA3870 | 879786 | -3.6164672 | 8.84488E-07 | 4.17433E-05 |
| narH | PA3874 | 879783 | -2.7438831 | 4.14872E-07 | 2.04462E-05 |
| narI | PA3872 | 878884 | -3.8055887 | 1.18407E-06 | 5.44968E-05 |
| narJ | PA3873 | 879782 | -3.684603 | 8.23638E-06 | 0.00033246 |
| nirN | PA0509 | 878428 | -2.0996904 | 9.61039E-08 | 5.29903E-06 |
| nosL | PA3396 | 879912 | -1.1866243 | 1.84851E-05 | 0.000681746 |

| | | | | | |
|---|---|---|---|---|---|
| nosY | PA3395 | 879978 | -2.2085383 | 5.23158E-06 | 0.000217423 |
| oprN | PA2495 | 882885 | -5.952706 | 0 | 0 |
| PA0510 | PA0510 | 881871 | -2.0645015 | 1.63289E-10 | 1.16584E-08 |
| PA1333 | PA1333 | 879337 | -2.6089482 | 9.48303E-27 | 1.20025E-24 |
| PA1416 | PA1416 | 881714 | -1.477368 | 1.08869E-07 | 5.7742E-06 |
| PA1417 | PA1417 | 881729 | -1.6373963 | 6.01451E-08 | 3.34948E-06 |
| PA1418 | PA1418 | 881711 | -1.0815197 | 0.000745889 | 0.016954506 |
| PA1419 | PA1419 | 881764 | -2.1351616 | 1.37659E-11 | 1.03598E-09 |
| PA1420 | PA1420 | 881752 | -1.9498341 | 1.35265E-16 | 1.29877E-14 |
| PA1743 | PA1743 | 881953 | -4.162325 | 0 | 0 |
| PA1744 | PA1744 | 881964 | -4.5559154 | 0 | 0 |
| PA1942 | PA1942 | 881774 | -5.9875665 | 0 | 0 |
| PA1970 | PA1970 | 880614 | -3.4925134 | 0 | 0 |
| PA2354 | PA2354 | 882192 | -1.1118798 | 5.08345E-05 | 0.001645916 |
| PA2358 | PA2358 | 878242 | -1.3258629 | 0.000991892 | 0.021920035 |
| PA2485 | PA2485 | 880511 | -3.494606 | 0 | 0 |
| PA2486 | PA2486 | 880512 | -4.5277753 | 0 | 0 |
| PA2487 | PA2487 | 882890 | -3.699953 | 0 | 0 |
| PA2491 | PA2491 | 880416 | -2.1825898 | 0 | 0 |
| PA2758 | PA2758 | 882947 | -1.5831227 | 4.72536E-24 | 5.48241E-22 |
| PA2759 | PA2759 | 882718 | -2.0236518 | 1.61882E-32 | 2.57578E-30 |
| PA2811 | PA2811 | 880412 | -1.345825 | 9.50273E-19 | 9.80013E-17 |
| PA2812 | PA2812 | 880277 | -1.212229 | 5.64635E-09 | 3.53309E-07 |
| PA2813 | PA2813 | 880278 | -2.5258262 | 9.4E-44 | 2.0833E-41 |
| PA3229 | PA3229 | 882400 | -6.1241274 | 0 | 0 |
| PA3230 | PA3230 | 882391 | -3.2442453 | 0 | 0 |
| PA3282 | PA3282 | 882445 | -1.5183458 | 9.22101E-06 | 0.000361632 |
| PA3283 | PA3283 | 882446 | -1.139733 | 2.07531E-06 | 9.1003E-05 |
| PA3381 | PA3381 | 878142 | -1.7253764 | 0.000188927 | 0.005260667 |
| PA3871 | PA3871 | 878883 | -3.84262 | 1.43919E-06 | 6.56954E-05 |
| PA4354 | PA4354 | 881441 | -3.829768 | 0 | 0 |
| PA4355 | PA4355 | 881442 | -2.8832977 | 7.28918E-32 | 1.09712E-29 |
| PA4623 | PA4623 | 881169 | -5.78145 | 0 | 0 |
| PA4881 | PA4881 | 882511 | -6.4857426 | 4.33985E-32 | 6.7135E-30 |
| PA4882 | PA4882 | 882512 | -1.9930938 | 4.33136E-37 | 7.53792E-35 |
| pagL | PA4661 | 881344 | -1.2285647 | 1.62769E-22 | 1.77738E-20 |
| xenB | PA4356 | 881382 | -2.39282 | 2.86183E-23 | 3.1875E-21 |
