## Supplemental Table 2 for "Modulation of the *Pseudomonas aeruginosa* quorum sensing cascade by MexT-regulated factors"

**Supplemental Table 2.** MexT regulon at an $OD_{600}$ of 2.0.

| Gene Symbol | Locus Tag | Gene ID | log2FC_MexT_OD2_WT_OD2 | pVal_MexT_OD2_WT_OD2 | pAdj_MexT_OD2_WT_OD2 |
|---|---|---|---|---|---|
| fpr | PA3397 | 879913 | 1.1045078 | 6.85534E-05 | 0.001186034 |
| PA0122 | PA0122 | 880611 | 2.8145795 | 0 | 0 |
| PA0130 | PA0130 | 880875 | 1.0989678 | 8.18348E-06 | 0.000179424 |
| PA0132 | PA0132 | 879350 | 1.6323024 | 5.28597E-05 | 0.000952672 |
| PA1656 | PA1656 | 882166 | 1.113103 | 1.30144E-06 | 3.37102E-05 |
| PA1657 | PA1657 | 879559 | 1.4086361 | 6.94796E-06 | 0.000157289 |
| PA1658 | PA1658 | 879592 | 1.4465269 | 7.29373E-06 | 0.000162475 |
| PA1659 | PA1659 | 880734 | 1.2742839 | 1.89831E-05 | 0.000381649 |
| PA1660 | PA1660 | 880603 | 1.1200874 | 6.91911E-08 | 2.18935E-06 |
| PA1664 | PA1664 | 880635 | 1.5847932 | 5.50696E-05 | 0.000979817 |
| PA1665 | PA1665 | 881944 | 1.6456726 | 6.58402E-05 | 0.001149418 |
| PA1666 | PA1666 | 882020 | 1.6189243 | 2.94475E-06 | 7.16127E-05 |
| PA1667 | PA1667 | 881962 | 1.7532798 | 1.12547E-11 | 5.01421E-10 |
| PA1668 | PA1668 | 882100 | 2.1305225 | 1.40734E-17 | 8.33773E-16 |
| PA1669 | PA1669 | 882077 | 1.7142956 | 3.33265E-18 | 2.06217E-16 |
| PA1869 | PA1869 | 881955 | 2.5914545 | 1.99544E-36 | 2.26787E-34 |
| PA2371 | PA2371 | 877948 | 1.0361449 | 1.49633E-08 | 4.98985E-07 |
| PA2763a | PA2763a | 17373368 | 1.0763892 | 9.00648E-05 | 0.00150171 |
| PA3326 | PA3326 | 882491 | 1.4495443 | 4.8419E-10 | 1.89891E-08 |
| PA3334 | PA3334 | 882499 | 4.016373 | 0 | 2.8E-45 |
| PA3521 | PA3521 | 878905 | 1.0042175 | 0.003222804 | 0.03199251 |
| PA3523 | PA3523 | 879790 | 1.6197994 | 2.56849E-07 | 7.5682E-06 |
| PA3574a | PA3574a | 17373374 | 2.0559936 | 4.91886E-07 | 1.36966E-05 |
| PA4506 | PA4506 | 881159 | 1.0300689 | 1.42917E-07 | 4.38335E-06 |
| PA4649 | PA4649 | 881286 | 1.0987594 | 1.69029E-05 | 0.000348638 |
| PA4650 | PA4650 | 881294 | 1.1732821 | 6.85739E-07 | 1.84487E-05 |
| PA4878 | PA4878 | 882234 | 1.2236124 | 1.311E-05 | 0.000276552 |
| PA5266 | PA5266 | 879578 | 1.1973057 | 8.82119E-05 | 0.001484145 |
| PA0050 | PA0050 | 878575 | 1.6996115 | 9.47531E-15 | 5.17333E-13 |
| phzH | PA0051 | 878637 | 1.457173 | 7.906E-12 | 3.57955E-10 |
| PA0112 | PA0112 | 879408 | 1.2184486 | 0.000816508 | 0.010357937 |
| PA0123 | PA0123 | 877571 | 1.3901081 | 1.59415E-21 | 1.18371E-19 |
| opdC | PA0162 | 879518 | 1.8529296 | 4.04902E-33 | 4.5098E-31 |
| PA0165 | PA0165 | 879418 | 1.1412905 | 1.18871E-09 | 4.44291E-08 |
| PA0174 | PA0174 | 877836 | 1.1020726 | 0.001013939 | 0.012410171 |
| PA0175 | PA0175 | 882282 | 1.4837809 | 0.003827669 | 0.036375914 |
| aer2 | PA0176 | 882219 | 1.1995925 | 0.005772472 | 0.049004413 |

| | | | | | |
|---|---|---|---|---|---|
| PA0187 | PA0187 | 882255 | 1.2646757 | 5.4862E-05 | 0.000979252 |
| hcpC | PA0263 | 880866 | 2.5057027 | 1.17082E-17 | 7.0111E-16 |
| metK | PA0546 | 878379 | 1.2136652 | 1.07684E-06 | 2.82875E-05 |
| PA0547 | PA0547 | 879420 | 1.023865 | 8.46374E-09 | 2.90954E-07 |
| cbpD | PA0852 | 877771 | 1.7645105 | 5.13596E-16 | 2.94867E-14 |
| oprD | PA0958 | 881970 | 1.5391793 | 1.817E-19 | 1.20463E-17 |
| rhlC | PA1130 | 877665 | 1.7842242 | 3.27692E-19 | 2.14696E-17 |
| PA1131 | PA1131 | 878079 | 1.1740103 | 7.40892E-10 | 2.82605E-08 |
| PA1211 | PA1211 | 882007 | 2.084553 | 3.43259E-05 | 0.000645814 |
| PA1212 | PA1212 | 882023 | 3.836752 | 6.20769E-12 | 2.83366E-10 |
| PA1213 | PA1213 | 882124 | 3.5433474 | 2.52811E-09 | 9.26251E-08 |
| PA1214 | PA1214 | 882040 | 2.937446 | 7.63602E-08 | 2.38904E-06 |
| PA1215 | PA1215 | 882005 | 2.9759848 | 4.26472E-09 | 1.51275E-07 |
| PA1216 | PA1216 | 878403 | 3.917679 | 2.19545E-15 | 1.21054E-13 |
| PA1217 | PA1217 | 879659 | 3.4835231 | 2.36276E-17 | 1.37065E-15 |
| PA1218 | PA1218 | 878233 | 3.9589984 | 3.37007E-21 | 2.37569E-19 |
| PA1219 | PA1219 | 880899 | 3.2660854 | 5.14143E-10 | 2.00228E-08 |
| PA1220 | PA1220 | 879715 | 3.5688412 | 3.83995E-09 | 1.37081E-07 |
| PA1221 | PA1221 | 878373 | 3.247769 | 3.40715E-30 | 3.38829E-28 |
| PA1301 | PA1301 | 881570 | 1.1751856 | 3.80916E-05 | 0.000709357 |
| cyoA | PA1317 | 881513 | 2.5046484 | 2.08345E-15 | 1.16027E-13 |
| cyoB | PA1318 | 881481 | 2.4227154 | 1.14928E-07 | 3.55574E-06 |
| cyoC | PA1319 | 881595 | 2.227335 | 2.08151E-05 | 0.000409608 |
| cyoD | PA1320 | 881604 | 1.9996376 | 0.000201791 | 0.003095804 |
| cyoE | PA1321 | 881575 | 1.7020034 | 0.000420089 | 0.005879904 |
| hcpA | PA1512 | 883081 | 2.4815075 | 7.59239E-18 | 4.64638E-16 |
| PA1521 | PA1521 | 883114 | 1.256622 | 5.71803E-05 | 0.001014131 |
| PA1550 | PA1550 | 883057 | 1.0271091 | 2.89269E-06 | 7.09664E-05 |
| PA1551 | PA1551 | 883008 | 1.0869861 | 3.83722E-07 | 1.09587E-05 |
| stp1 | PA1670 | 882277 | 1.6718878 | 1.73855E-11 | 7.68413E-10 |
| stk1 | PA1671 | 883070 | 1.1093968 | 6.03482E-08 | 1.93149E-06 |
| pscO | PA1696 | 878091 | 1.5575796 | 1.39195E-05 | 0.000292519 |
| PA1697 | PA1697 | 879763 | 1.1544354 | 6.93757E-06 | 0.000157289 |
| pscB | PA1715 | 879568 | 1.2806069 | 0.004848334 | 0.043339282 |
| PA1854 | PA1854 | 882032 | 1.0284944 | 0.001200154 | 0.014173113 |
| PA1855 | PA1855 | 880682 | 1.6997409 | 1.99087E-05 | 0.000394561 |
| PA1870 | PA1870 | 877596 | 1.1114173 | 1.66057E-05 | 0.000345063 |
| lasA | PA1871 | 878260 | 1.7627566 | 6.90027E-16 | 3.92118E-14 |
| phzA2 | PA1899 | 878477 | 5.625596 | 0 | 0 |
| phzB2 | PA1900 | 878211 | 5.890583 | 0 | 0 |

| | | | | | |
|---|---|---|---|---|---|
| phzC2 | PA1901 | 882061 | 4.106911 | 0 | 0 |
| phzD2 | PA1902 | 881873 | 5.40953 | 0 | 0 |
| phzE2 | PA1903 | 882029 | 5.047485 | 0 | 0 |
| phzF2 | PA1904 | 879183 | 5.0867105 | 0 | 0 |
| phzG2 | PA1905 | 880482 | 4.8029146 | 0 | 0 |
| PA1906 | PA1906 | 881995 | 2.8461294 | 1.07339E-22 | 8.07795E-21 |
| PA1907 | PA1907 | 879184 | 2.2237427 | 6.02225E-09 | 2.1093E-07 |
| PA1914 | PA1914 | 878271 | 1.7490927 | 9.42346E-13 | 4.52407E-11 |
| pcoB | PA2064 | 877759 | 1.7687854 | 3.9523E-08 | 1.27967E-06 |
| pcoA | PA2065 | 877718 | 1.5574735 | 1.75954E-11 | 7.71565E-10 |
| PA2066 | PA2066 | 877703 | 4.52587 | 0 | 0 |
| PA2067 | PA2067 | 878262 | 3.9280777 | 5.44433E-32 | 5.83067E-30 |
| PA2068 | PA2068 | 877702 | 5.6560087 | 0 | 0 |
| PA2069 | PA2069 | 877951 | 6.121604 | 0 | 0 |
| hcnA | PA2193 | 882197 | 1.4895806 | 8.82186E-11 | 3.72189E-09 |
| hcnB | PA2194 | 882182 | 1.8370924 | 7.66803E-25 | 6.27989E-23 |
| hcnC | PA2195 | 882196 | 1.855879 | 1.73641E-13 | 8.79095E-12 |
| PA2196 | PA2196 | 881967 | 1.0343953 | 2.39573E-07 | 7.09671E-06 |
| PA2274 | PA2274 | 882293 | 1.947892 | 1.97113E-09 | 7.2697E-08 |
| chiC | PA2300 | 878587 | 3.1345396 | 2.81169E-40 | 3.40397E-38 |
| PA2327 | PA2327 | 883075 | 1.101008 | 2.49667E-08 | 8.17881E-07 |
| PA2328 | PA2328 | 881900 | 1.2203404 | 8.76134E-13 | 4.24277E-11 |
| PA2330 | PA2330 | 883026 | 1.133087 | 9.4157E-10 | 3.54298E-08 |
| PA2331 | PA2331 | 881926 | 1.4184358 | 5.05738E-21 | 3.52057E-19 |
| PA2448 | PA2448 | 882921 | 1.0341737 | 2.88629E-10 | 1.15639E-08 |
| xylY | PA2517 | 882902 | 1.2014062 | 0.000648693 | 0.008601356 |
| PA2564 | PA2564 | 882722 | 1.4372817 | 2.13634E-14 | 1.14397E-12 |
| PA2565 | PA2565 | 882723 | 1.8989971 | 7.7416E-14 | 3.99194E-12 |
| PA2566 | PA2566 | 882794 | 1.2896307 | 2.26822E-20 | 1.55947E-18 |
| lecA | PA2570 | 882335 | 3.140037 | 0 | 6.2E-44 |
| PA2788 | PA2788 | 882783 | 1.6979694 | 6.77218E-10 | 2.60098E-08 |
| atuG | PA2892 | 882587 | 1.3179512 | 0.000226395 | 0.003416788 |
| pelG | PA3058 | 879929 | 1.2538701 | 5.1461E-13 | 2.53616E-11 |
| pelF | PA3059 | 879152 | 1.4195709 | 1.77768E-13 | 8.91881E-12 |
| pelD | PA3061 | 880405 | 1.4552052 | 3.21273E-09 | 1.1618E-07 |
| PA3325 | PA3325 | 882490 | 1.1521326 | 8.06755E-12 | 3.62324E-10 |
| PA3327 | PA3327 | 882492 | 1.7772273 | 9.28633E-11 | 3.88839E-09 |
| PA3328 | PA3328 | 882493 | 2.2678761 | 3.25028E-23 | 2.47956E-21 |
| PA3329 | PA3329 | 882494 | 2.7147896 | 0 | 0 |
| PA3330 | PA3330 | 882495 | 3.7626033 | 0 | 5.2E-44 |

| | | | | | |
|---|---|---|---|---|---|
| PA3331 | PA3331 | 882496 | 3.9879947 | 5.16E-43 | 6.6719E-41 |
| PA3332 | PA3332 | 882497 | 4.45628 | 0 | 0 |
| fabH2 | PA3333 | 882498 | 3.999765 | 1.2584E-41 | 1.55735E-39 |
| PA3335 | PA3335 | 882500 | 3.7590766 | 1.12533E-36 | 1.30561E-34 |
| PA3336 | PA3336 | 882501 | 2.0141146 | 2.46359E-06 | 6.09765E-05 |
| lecB | PA3361 | 882528 | 3.3477588 | 0 | 0 |
| rhlI | PA3476 | 878967 | 1.1454918 | 5.08834E-12 | 2.3419E-10 |
| rhlB | PA3478 | 878954 | 3.2005177 | 0 | 0 |
| rhlA | PA3479 | 878955 | 4.1239204 | 0 | 0 |
| PA3516 | PA3516 | 879993 | 1.0295094 | 0.000484433 | 0.006564008 |
| PA3517 | PA3517 | 879063 | 1.0512273 | 0.003658171 | 0.035070404 |
| PA3519 | PA3519 | 878876 | 1.9477133 | 8.03172E-05 | 0.001363679 |
| PA3520 | PA3520 | 878877 | 2.385776 | 6.42307E-14 | 3.37454E-12 |
| adk | PA3686 | 879082 | 1.1884669 | 3.30441E-11 | 1.40475E-09 |
| lasB | PA3724 | 880368 | 2.1654446 | 2.70765E-25 | 2.28469E-23 |
| PA3727 | PA3727 | 880348 | 1.0260646 | 1.76818E-06 | 4.45565E-05 |
| PA3734 | PA3734 | 880327 | 1.2263174 | 2.36392E-13 | 1.17542E-11 |
| narK1 | PA3877 | 878320 | 2.2162707 | 0.000221462 | 0.003360554 |
| PA3892 | PA3892 | 878746 | 1.2633095 | 2.30159E-07 | 6.8543E-06 |
| PA3893 | PA3893 | 878747 | 1.3935153 | 2.60255E-09 | 9.47293E-08 |
| PA3894 | PA3894 | 878791 | 1.0726025 | 1.31422E-06 | 3.38838E-05 |
| PA3920 | PA3920 | 879022 | 1.6655331 | 0.000431401 | 0.006021227 |
| cioA | PA3930 | 879045 | 1.2573134 | 4.71924E-24 | 3.75449E-22 |
| ladS | PA3974 | 878890 | 1.5078326 | 7.03126E-33 | 7.67786E-31 |
| PA3990 | PA3990 | 878937 | 1.0239191 | 0.000457167 | 0.006317529 |
| hpcG | PA4127 | 880260 | 1.1116729 | 8.69985E-13 | 4.24277E-11 |
| PA4130 | PA4130 | 881847 | 1.1720622 | 2.09053E-21 | 1.53186E-19 |
| PA4131 | PA4131 | 881848 | 1.9910942 | 0 | 0 |
| PA4132 | PA4132 | 880050 | 1.5919966 | 6.35659E-20 | 4.31706E-18 |
| PA4133 | PA4133 | 880051 | 2.6146755 | 4.11986E-29 | 3.82392E-27 |
| PA4134 | PA4134 | 880084 | 2.2471962 | 1.91509E-29 | 1.80765E-27 |
| PA4136 | PA4136 | 880090 | 1.049065 | 2.14184E-08 | 7.05792E-07 |
| PA4141 | PA4141 | 880096 | 3.0694606 | 0 | 0 |
| PA4142 | PA4142 | 880112 | 1.6154714 | 1.51235E-17 | 8.86554E-16 |
| PA4143 | PA4143 | 880113 | 1.4100789 | 2.07067E-15 | 1.16027E-13 |
| PA4144 | PA4144 | 880262 | 1.4455621 | 5.82568E-07 | 1.6061E-05 |
| mexG | PA4205 | 880449 | 1.9883562 | 1.0568E-30 | 1.07006E-28 |
| mexH | PA4206 | 880450 | 1.9605036 | 1.0018E-17 | 6.06417E-16 |
| mexI | PA4207 | 880346 | 1.8083084 | 4.45901E-28 | 4.0052E-26 |
| opmD | PA4208 | 880347 | 2.425414 | 2.4721E-18 | 1.54686E-16 |

| | | | | | |
|---|---|---|---|---|---|
| phzM | PA4209 | 880514 | 3.6457636 | 0 | 0 |
| phzA1 | PA4210 | 880515 | 4.7093506 | 6.26358E-24 | 4.91294E-22 |
| phzB1 | PA4211 | 880518 | 6.0325694 | 0 | 0 |
| phzC1 | PA4212 | 880519 | 4.0002785 | 1.42519E-08 | 4.78127E-07 |
| phzG1 | PA4216 | 881835 | 5.288512 | 1.04333E-29 | 1.00178E-27 |
| phzS | PA4217 | 881836 | 5.2363553 | 0 | 0 |
| PA4223 | PA4223 | 880074 | 1.4883937 | 0.00358151 | 0.03480877 |
| pchG | PA4224 | 880082 | 2.0340276 | 0.001221885 | 0.014325632 |
| pchF | PA4225 | 880083 | 1.6786491 | 2.14007E-05 | 0.000419649 |
| pchB | PA4230 | 881846 | 1.4665633 | 3.54958E-07 | 1.03495E-05 |
| pchA | PA4231 | 881821 | 1.6205597 | 1.81569E-11 | 7.89966E-10 |
| PA4384 | PA4384 | 881389 | 1.0583689 | 7.10921E-08 | 2.23679E-06 |
| PA4469 | PA4469 | 881041 | 1.0102013 | 0.000204113 | 0.003122818 |
| PA4471 | PA4471 | 881034 | 1.1421306 | 0.000125383 | 0.002018084 |
| phuR | PA4710 | 881550 | 1.1756346 | 0.005235258 | 0.046065245 |
| phaC2 | PA5058 | 881185 | 1.0985643 | 2.05779E-18 | 1.30225E-16 |
| rmlB | PA5161 | 879982 | 1.0630944 | 8.23219E-19 | 5.33082E-17 |
| PA5219 | PA5219 | 882291 | 2.2050803 | 0 | 0 |
| PA5220 | PA5220 | 882292 | 3.2145371 | 3.359E-42 | 4.2506E-40 |
| hcpB | PA5267 | 879579 | 2.2252288 | 1.89834E-12 | 8.9592E-11 |
| atpB | PA5560 | 880089 | 1.0933278 | 1.75694E-07 | 5.31759E-06 |
| PA0078 | PA0078 | 879392 | -1.0359347 | 0.001274876 | 0.014822099 |
| PA0510 | PA0510 | 881871 | -1.378441 | 1.77436E-05 | 0.000364628 |
| PA1978 | PA1978 | 880879 | -1.3936888 | 2.86322E-05 | 0.000544208 |
| PA2486 | PA2486 | 880512 | -4.7765036 | 0 | 0 |
| PA2491 | PA2491 | 880416 | -1.1868284 | 1.36667E-19 | 9.16986E-18 |
| PA4355 | PA4355 | 881442 | -2.6111999 | 4.53394E-25 | 3.76858E-23 |
| PA5168 | PA5168 | 877772 | -1.0510042 | 0.000060637 | 0.001072024 |
| bioF | PA0501 | 879619 | -1.2430855 | 2.26876E-07 | 6.79286E-06 |
| bioD | PA0504 | 878220 | -1.1266121 | 1.64371E-07 | 5.00208E-06 |
| nirN | PA0509 | 878428 | -1.6958036 | 1.59958E-05 | 0.000333905 |
| PA0839 | PA0839 | 882078 | -1.702509 | 7.5522E-10 | 2.8611E-08 |
| PA1190 | PA1190 | 880589 | -1.0800213 | 3.66181E-05 | 0.000684316 |
| PA1231 | PA1231 | 881164 | -1.4084458 | 0.001979433 | 0.021404782 |
| PA1333 | PA1333 | 879337 | -2.7799306 | 3.50017E-30 | 3.41973E-28 |
| PA1743 | PA1743 | 881953 | -4.038801 | 0 | 0 |
| PA1744 | PA1744 | 881964 | -4.085281 | 0 | 0 |
| PA1942 | PA1942 | 881774 | -6.575373 | 0 | 0 |
| PA1970 | PA1970 | 880614 | -4.827661 | 0 | 0 |
| PA1975 | PA1975 | 877962 | -2.187948 | 1.44509E-14 | 7.81329E-13 |

| | | | | | |
|---|---|---|---|---|---|
| ercS' | PA1976 | 880111 | -2.3165855 | 2.58364E-21 | 1.84466E-19 |
| PA1977 | PA1977 | 878077 | -2.7406213 | 8.08556E-06 | 0.000178684 |
| eraS | PA1979 | 880447 | -2.480119 | 6.73451E-14 | 3.50509E-12 |
| eraR | PA1980 | 880513 | -2.6989477 | 2.87489E-12 | 1.3454E-10 |
| PA1981 | PA1981 | 880196 | -2.813988 | 0.00079026 | 0.01009376 |
| exaA | PA1982 | 880475 | -3.0545344 | 1.94243E-24 | 1.56773E-22 |
| exaB | PA1983 | 880387 | -2.9416 | 3.74479E-08 | 1.21958E-06 |
| exaC | PA1984 | 880413 | -1.0172932 | 4.7382E-09 | 1.67007E-07 |
| pqqB | PA1986 | 880354 | -1.3043365 | 6.57506E-07 | 1.78617E-05 |
| pqqC | PA1987 | 879112 | -1.376477 | 3.62932E-07 | 1.04724E-05 |
| pqqD | PA1988 | 879085 | -1.3795372 | 3.29576E-10 | 1.30171E-08 |
| pqqE | PA1989 | 880128 | -1.1139989 | 0.000692978 | 0.009080458 |
| pqqH | PA1990 | 880121 | -1.1840495 | 2.50144E-05 | 0.000480363 |
| bdhA | PA2003 | 879996 | -1.0545119 | 2.01664E-06 | 5.0137E-05 |
| PA2026 | PA2026 | 877995 | -1.2798847 | 9.71838E-11 | 4.03893E-09 |
| PA2317 | PA2317 | 883103 | -1.1475292 | 3.73331E-06 | 8.84716E-05 |
| gntR | PA2320 | 883124 | -1.3597336 | 1.91203E-06 | 4.79644E-05 |
| PA2338 | PA2338 | 883086 | -1.1340393 | 1.48373E-10 | 6.0313E-09 |
| PA2341 | PA2341 | 878622 | -1.056754 | 3.08501E-05 | 0.000582388 |
| bexR | PA2432 | 882852 | -1.1054603 | 8.01894E-14 | 4.09702E-12 |
| PA2482 | PA2482 | 879889 | -1.0396509 | 1.21584E-05 | 0.000257452 |
| PA2485 | PA2485 | 880511 | -4.480828 | 0 | 0 |
| PA2487 | PA2487 | 882890 | -3.672121 | 0 | 0 |
| mexT | PA2492 | 880417 | -6.845947 | 0 | 0 |
| mexE | PA2493 | 880212 | -7.1676016 | 0 | 0 |
| mexF | PA2494 | 882884 | -7.0272865 | 0 | 0 |
| oprN | PA2495 | 882885 | -6.8770933 | 0 | 0 |
| PA2758 | PA2758 | 882947 | -1.6863893 | 9.02283E-28 | 7.85127E-26 |
| PA2759 | PA2759 | 882718 | -3.016338 | 0 | 0 |
| PA2811 | PA2811 | 880412 | -1.3289112 | 1.83524E-18 | 1.17476E-16 |
| PA2812 | PA2812 | 880277 | -1.2920494 | 5.1801E-10 | 2.00333E-08 |
| PA2813 | PA2813 | 880278 | -2.0968688 | 6.49122E-31 | 6.69437E-29 |
| PA2814 | PA2814 | 882804 | -1.6401517 | 1.01757E-10 | 4.19766E-09 |
| PA2833 | PA2833 | 882817 | -1.1537558 | 0.000606077 | 0.008055472 |
| PA3079 | PA3079 | 882959 | -1.0846925 | 2.23998E-05 | 0.00043617 |
| csaA | PA3221 | 882552 | -1.0058683 | 2.61368E-05 | 0.000500191 |
| PA3229 | PA3229 | 882400 | -6.6184177 | 0 | 0 |
| PA3230 | PA3230 | 882391 | -3.8083062 | 0 | 0 |
| PA3231 | PA3231 | 882392 | -1.4323058 | 3.06362E-05 | 0.000580317 |
| PA3282 | PA3282 | 882445 | -1.1329569 | 0.001086626 | 0.013241622 |

| | | | | | |
|---|---|---|---|---|---|
| PA3283 | PA3283 | 882446 | -1.0243664 | 2.04672E-05 | 0.000404192 |
| amiC | PA3364 | 877798 | -1.0002799 | 6.47474E-05 | 0.001133894 |
| PA3365 | PA3365 | 877799 | -1.4626061 | 1.91331E-10 | 7.72116E-09 |
| nosL | PA3396 | 879912 | -1.7979099 | 1.03247E-10 | 4.22783E-09 |
| PA3432 | PA3432 | 878984 | -1.1305162 | 0.000281667 | 0.004117063 |
| PA3511 | PA3511 | 879160 | -1.1956878 | 0.000192408 | 0.002968198 |
| arnE | PA3557 | 879132 | -1.124755 | 0.000316415 | 0.004565064 |
| PA3568 | PA3568 | 878696 | -1.4252354 | 1.07437E-06 | 2.82875E-05 |
| mmsB | PA3569 | 879097 | -1.0338805 | 3.9626E-05 | 0.000733147 |
| PA3602 | PA3602 | 880138 | -1.1207148 | 2.15937E-23 | 1.67021E-21 |
| PA3709 | PA3709 | 880410 | -1.2849684 | 7.3621E-07 | 1.9617E-05 |
| PA3973 | PA3973 | 878595 | -1.4615314 | 0.001782441 | 0.019529654 |
| PA4149 | PA4149 | 880109 | -1.0872767 | 1.8217E-05 | 0.000370616 |
| fepG | PA4161 | 880214 | -1.5039257 | 1.10453E-06 | 2.87435E-05 |
| PA4353 | PA4353 | 881435 | -1.5042355 | 5.39637E-14 | 2.86213E-12 |
| PA4354 | PA4354 | 881441 | -3.69468 | 0 | 0 |
| xenB | PA4356 | 881382 | -2.83294 | 6.1345E-32 | 6.44585E-30 |
| PA4623 | PA4623 | 881169 | -6.717972 | 0 | 0 |
| pagL | PA4661 | 881344 | -1.3771894 | 7.08431E-28 | 6.2623E-26 |
| PA4881 | PA4881 | 882511 | -7.247754 | 1.11096E-39 | 1.31637E-37 |
| PA4980 | PA4980 | 880426 | -1.0614209 | 2.78689E-05 | 0.000531513 |
| PA5097 | PA5097 | 878367 | -1.896876 | 2.28612E-21 | 1.65343E-19 |
| hutH | PA5098 | 878368 | -2.210363 | 2.07018E-28 | 1.88998E-26 |
| PA5099 | PA5099 | 880889 | -2.3486679 | 0 | 0 |
| hutU | PA5100 | 880890 | -1.5749017 | 8.00028E-27 | 6.8544E-25 |
| PA5144 | PA5144 | 878777 | -1.3558519 | 0.003254024 | 0.032130603 |
| PA5182 | PA5182 | 881732 | -1.0018505 | 0.000571571 | 0.007633286 |
| glcE | PA5354 | 877711 | -1.1312255 | 8.57476E-05 | 0.001447056 |
