## Supplemental Table 3 for "Modulation of the *Pseudomonas aeruginosa* quorum sensing cascade by MexT-regulated factors"

**Supplemental Table 3. Bacterial strains used in this study.**

| Bacterial strain | Description | Source |
|---|---|---|
| ***P. aeruginosa*** | | |
| PAO1 | Wild-type strain | (1) |
| PAO1Δ*lasR* | PAO1 with unmarked, in-frame *lasR* deletion | (2) |
| PAO1Δ*lasR* Δ*mexT* | PAO1 with unmarked, in-frame *lasR* and *mexT* deletion | This work |
| PAO1Δ*mexT* Δ*pqsE* | PAO1 with unmarked, in-frame *mexT* and *pqsE* deletion | (3) |
| PAO1Δ*mexT* | PAO1 with unmarked, in-frame *mexT* deletion | (3) |
| PAO1Δ*mexEF* | PAO1 with unmarked, in-frame *mexEF* deletion | This work |
| PAO1 attn7::$P_{araBAD}$-*pqsE* | Made by introducing pUC18-miniTn7T-*araBAD pqsE* to PAO1 | This work |
| PAO1 attn7::$P_{araBAD}$-*mexEF-oprN* | Made by introducing pUC18-miniTn7T-*araBAD mexEF-oprN* to PAO1 | This work |
| E192 | Cystic fibrosis lung clinical *P. aeruginosa* isolate | (4) |
| E192 Δ*lasR* | E192 with unmarked, in-frame *lasR* deletion | This work |
| E192 Δ*mexT* | E192 with unmarked, in-frame *mexT* deletion | This work |
| E192 Δ*lasR* Δ*mexT* | E192 with unmarked, in-frame *lasR* and *mexT* deletion | This work |
| E194 | Cystic fibrosis lung clinical *P. aeruginosa* isolate | (4) |
| E194 Δ*lasR* | E194 with unmarked, in-frame *lasR* deletion | This work |
| E194 Δ*mexT* | E194 with unmarked, in-frame *mexT* deletion | This work |
| E194 Δ*lasR* Δ*mexT* | E194 with unmarked, in-frame *lasR* and *mexT* deletion | This work |
| E195 | Cystic fibrosis lung clinical *P. aeruginosa* isolate | (4) |
| E195 Δ*lasR* | E195 with unmarked, in-frame *lasR* deletion | This work |
| E195 Δ*mexT* | E195 with unmarked, in-frame *mexT* deletion | This work |
| E195 Δ*lasR* Δ*mexT* | E195 with unmarked, in-frame *lasR* and *mexT* deletion | This work |
| ***E. coli*** | | |
| NEB5α | *fhuA2Δ(argF-lacZ)U169 phoA glnV44 Φ80Δ(lacZ)M15 gyrA96 recA1 relA1* | NEB |
| S17-1 | *recA pro hsdR RP4-2Tc::Mu-Km::Tn7* | (5) |
