## Supplemental Table 4 for "Modulation of the *Pseudomonas aeruginosa* quorum sensing cascade by MexT-regulated factors"

**Supplemental Table 4. Plasmids used in this study.**

| Plasmids | Description | Source |
|---|---|---|
| pEXG2 | Allelic exchange vector with pBR origin, *sacB*, $Gm^{R*}$ | (6) |
| pEXG2-PAO1-*mexT*-KO | pEXG2 containing sequences for PAO1 *mexT* in-frame deletion | (7) |
| pEXG2-PAO1-*mexEF*-KO | pEXG2 containing sequences for PAO1 *mexEF* in-frame deletion | This work |
| pEXG2-E192-*lasR*-KO | pEXG2 containing sequences for E192 *lasR* in-frame deletion | This work |
| pEXG2-E192-*mexT*-KO | pEXG2 containing sequences for E192 *mexT* in-frame deletion | This work |
| pEXG2-E194-*lasR*-KO | pEXG2 containing sequences for E194 *lasR* in-frame deletion | This work |
| pEXG2-E194-*mexT*-KO | pEXG2 containing sequences for E194 *mexT* in-frame deletion | This work |
| pEXG2-E195-*lasR*-KO | pEXG2 containing sequences for E195 *lasR* in-frame deletion | This work |
| pEXG2-E195-*mexT*-KO | pEXG2 containing sequences for E195 *mexT* in-frame deletion | This work |
| pBBR1MCS-5 | Broad-host-range expression plasmid, $Gm^R$ | (8) |
| $pP_{lasI}$-gfp | pBBR1MCS-5 with *lasI* promoter fused to *gfp*, $Gm^R$; encodes -282 to +223 relative to the start of *lasI* and includes the complete *rsaL* binding site | (3) |
| $pP_{rhlA}$-gfp | pBBR1MCS-5 with *rhlA* promoter fused to *gfp*, $Gm^R$; encodes -500 to +31 relative to the start of *rhlA* | (3) |
| $pP_{pqsA}$-gfp | pBBR1MCS-5 with *pqsA* promoter fused to *gfp*, $Gm^R$; encodes -429 to +3 relative to the start of *pqsA* | (9) |
| $pP_{mexE}$-gfp | pBBR1MCS-5 with *mexE* promoter fused to *gfp*, $Gm^R$; encodes -500 to +31 relative to the start *mexE* | (3) |
| pUC18-mini-tn7T-*araBAD* | Suicide delivery vector with insertion at attTn7 site, $Gm^R$ | This study |
| pTNS2 | T7 transposase expression vector; R6K *ori*, *ori* T, $Amp^{R**}$ | (10) |
| pFLP2 | Site-specific excision vector; *sacB*, *ori* T, $Cb^{R***}$ | (11) |
| pUC18-mini-tn7T-*araBAD pqsE* | pUC18-mini-Tn7T-*araBAD* with $P_{araBAD}$-driven *pqsE* | This study |
| pUC18-mini-tn7T-*araBAD mexEF-oprN* | pUC18-mini-Tn7T-*araBAD* with $P_{araBAD}$-driven *mexEF-oprN* | This study |

* $Gm^R$, resistant to gentamicin

** $Amp^R$, resistant to ampicillin

*** $Cb^R$, resistant to carbenicillin

Supplemental references


1. Stover CK, Pham XQ, Erwin AL, Mizoguchi SD, Warrener P, Hickey MJ, Brinkman FS, Hufnagle WO, Kowalik DJ, Lagrou M, Garber RL, Goltry L, Tolentino E, Westbrock-Wadman S, Yuan Y, Brody LL, Coulter SN, Folger KR, Kas A, Larbig K, Lim R, Smith K, Spencer D, Wong GK, Wu Z, Paulsen IT, Reizer J, Saier MH, Hancock RE, Lory S, Olson MV. 2000. Complete genome sequence of *Pseudomonas aeruginosa* PAO1, an opportunistic pathogen. Nature 406:959-64.
2. Wang M, Schaefer AL, Dandekar AA, Greenberg EP. 2015. Quorum sensing and policing of *Pseudomonas aeruginosa* social cheaters. Proc Natl Acad Sci U S A 112:2187-91.
3. Kostylev M, Smalley NE, Chao MH, Greenberg EP. 2023. Relationship of the transcription factor MexT to quorum sensing and virulence in *Pseudomonas aeruginosa*. J Bacteriol 205:e0022623.
4. Feltner JB, Wolter DJ, Pope CE, Groleau MC, Smalley NE, Greenberg EP, Mayer-Hamblett N, Burns J, Deziel E, Hoffman LR, Dandekar AA. 2016. LasR variant cystic fibrosis isolates reveal an adaptable quorum-Sensing hierarchy in *Pseudomonas aeruginosa*. mBio 7.
5. Simon R, Priefer U, Puhler A. 1983. A broad host range mobilization system for *in vivo* genetic-engineering -- transposon mutagenesis in Gram-negative bacteria. Bio-Technology 1:784-791.
6. Rietsch A, Vallet-Gely I, Dove SL, Mekalanos JJ. 2005. ExsE, a secreted regulator of type III secretion genes in *Pseudomonas aeruginosa*. Proc Natl Acad Sci U S A 102:8006-11.
7. Kostylev M, Kim DY, Smalley NE, Salukhe I, Greenberg EP, Dandekar AA. 2019. Evolution of the *Pseudomonas aeruginosa* quorum-sensing hierarchy. Proc Natl Acad Sci U S A 116:7027-7032.
8. Kovach ME, Elzer PH, Hill DS, Robertson GT, Farris MA, Roop RM, 2nd, Peterson KM. 1995. Four new derivatives of the broad-host-range cloning vector pBBR1MCS, carrying different antibiotic-resistance cassettes. Gene 166:175-6.
9. Smalley NE, Schaefer AL, Asfahl KL, Perez C, Greenberg EP, Dandekar AA. 2022. Evolution of the quorum sensing regulon in cooperating populations of *Pseudomonas aeruginosa*. mBio 13:e0016122.
10. Choi KH, Gaynor JB, White KG, Lopez C, Bosio CM, Karkhoff-Schweizer RR, Schweizer HP. 2005. A Tn7-based broad-range bacterial cloning and expression system. Nat Methods 2:443-8.
11. Hoang TT, Karkhoff-Schweizer RR, Kutchma AJ, Schweizer HP. 1998. A broad-host-range Flp-FRT recombination system for site-specific excision of chromosomally-located DNA sequences: application for isolation of unmarked *Pseudomonas aeruginosa* mutants. Gene 212:77-86.
